# A novel Pyk2-derived peptide inhibits invadopodia-mediated breast cancer metastasis

**DOI:** 10.1101/2022.04.06.487297

**Authors:** Shams Twafra, Chana G. Sokolik, Tal Sneh, Kolluru D. Srikanth, Tomer Meirson, Alessandro Genna, Jordan H. Chill, Hava Gil-Henn

## Abstract

The non-receptor tyrosine kinase Pyk2 is highly expressed in breast cancer, where it mediates invadopodia formation and function via interaction with the actin-nucleation promoting factor cortactin. Here, we designed a cell-permeable peptide inhibitor that contains the second proline-rich region (PRR2) sequence of Pyk2, which binds to the SH3 domain of cortactin and blocks spontaneous lung metastasis in immune-competent mice by inhibiting invadopodia maturation and function. The native structure of the Pyk2-PRR2:cortactin-SH3 complex was determined using nuclear magnetic resonance (NMR), revealing an extended class II interaction surface spanning the canonical binding groove and a second hydrophobic surface which significantly contributes to ligand affinity. Using structure-guided design, we created a mutant peptide lacking critical residues involved in binding that failed to inhibit invadopodia maturation and function and consequent metastatic dissemination in mice. Our findings shed light on the specific molecular interactions between Pyk2 and cortactin and suggest that their inhibition may be used as a novel strategy for blocking breast cancer metastasis.

## INTRODUCTION

Metastatic dissemination of cancer cells from the primary tumor and their spread to distant tissues and organs is the leading cause of mortality in breast cancer patients. Although cytotoxic drugs and hormone-blocking therapeutics are often used to shrink the primary tumor or prevent disease recurrence, no treatment to permanently eradicate metastatic breast cancer exists at present. Metastatic cancer cells leaving the primary tumor use invasive feet-like structures called invadopodia, actin-rich protrusions with extracellular matrix (ECM) degrading activity, to invade through the basement membrane and to intravasate into blood vessels and spread to distant tissues and organs throughout the body. Invadopodia were identified in several invasive cancer cell lines, such as breast, head and neck, prostate, fibrosarcoma, and melanoma [1] as well as in primary tumor cells [2-4]. Moreover, direct molecular links between invadopodia assembly and cancer metastasis have previously been demonstrated in both mice models and human patients [5-7]. Along these lines, we have recently shown that therapeutic inhibition of invadopodia formation and function can block breast cancer metastasis in a xenograft mouse model [8].

The actin-nucleation promoting factor cortactin is a crucial regulator of actin cytoskeleton remodeling and an essential component of invadopodia. Cortactin plays an imperative role in actin polymerization-mediated membrane protrusions, cell migration, cell-cell adhesion, endocytosis, and neuronal synapse stabilization [9]. Its five functional regions include an N-terminal acidic domain (NTA) that binds the Arp3 component of the Arp2/3 complex and synergizes with the N-WASP VCA domain to promote actin polymerization, six and a half F-actin-binding repeats of 37 amino acids each, an α-helical domain with unknown function, a proline-serine-threonine (PST) rich phosphorylation region enabling cortactin to integrate signals from diverse upstream signaling cascades, and a Src homology 3 (SH3) domain (residues 495-542) [10]. The SH3 domain mediates the association of cortactin with various signaling proteins such as dynamin, Wiskott-Aldrich Syndrome (WASp)-interacting protein (WIP), CortBP-1/Shank2, Shank-3, ZO-1, and faciogenital dysplasia protein (Fgd1) [11-15]. The association of cortactin with proline-rich regions of diverse proteins enables regulating diverse actin remodeling-mediated cellular processes, such as endocytosis, lamellipodial protrusions, and cell migration [16]. Amplification of the *CTTN* gene followed by overexpression of cortactin have been observed in various human cancers and are associated with poor patient prognosis [8, 17, 18]. Importantly, cortactin is an essential component of invadopodia, regulating both their assembly and the trafficking and secretion of matrix metalloproteinases (MMPs) from invadopodia tips [19-21]. Cortactin phosphorylation on tyrosine residues regulates an on-off switch which initiates actin polymerization in invadopodia, leading to their maturation and activation and consequent tumor cell invasion [22].

Complexes between SH3 domains and proline-rich polypeptide segments are well-known regulators of major intracellular signaling events [23, 24]. A key feature is the steric compatibility between a flat SH3 hydrophobic surface with three grooves delineated by conserved aromatic residues and a PxxP core adopting a left-handed helical conformation known as the polyproline II helix [25, 26], recently also termed κ-helix [27]. Typically, two qP dipeptides (q being a hydrophobic residue) fill binding pockets between the conserved aromatic residues [28-30], and the inherent symmetry of the PxxP helix allows for binding in both possible orientations. The location of flanking basic residues and their interaction with negatively charged residues in the SH3 specificity pocket determines the preferred binding pose, roughly dividing SH3 ligands into class I and class II motifs, exhibiting N-terminal or C-terminal basic residues, respectively [28, 29, 31-33]. Screening of a phage-displayed peptide library defined the cortactin-binding consensus motif as +**PP**Ψ**P**x**KP** (‘Ψ’ is an aliphatic residue, >90% conserved positions in bold) [10, 34, 35], a class II motif with an additional PxxP motif, sometimes termed a class VIII motif [36]. However, structure determination of SH3/PxxP complexes is highly challenging due to their moderate affinity levels, which are in the micromolar range. Limited information at atomic resolution restricts a deeper structural understanding of cortactin SH3-PRR regulatory complexes.

Proline-rich tyrosine kinase 2 (Pyk2), along with focal adhesion kinase (FAK), comprise a subfamily of non-receptor protein tyrosine kinases that integrate signals from various receptors, activating signaling pathways that regulate proliferation, migration, and invasion in numerous cell types [37-39]. Both share a conserved domain structure, consisting of an N-terminal FERM domain, a central catalytic kinase domain, three proline-rich sequences, and a C-terminal FAT domain [40]. Two of the proline-rich sequences, proline-rich region 2 (^713^Pro-Pro-Pro-Lys Pro-Ser-Arg-Pro^720^) (PRR2) and proline-rich region 3 (^855^Pro-Pro-Gln-Lys-Pro-Pro-Arg-Leu^862^) (PRR3) [41] are major sites for SH3-mediated protein-protein interactions [42, 43]. The former exhibits high similarity between the two kinases, while the latter shows very little similarity, possibly accounting for their unique binding partners. Interestingly, a previous study showed that cortactin SH3 binds directly to both FAK-PRR2 and FAK-PRR3, an interaction followed by a sequence of events that promote focal adhesion turnover and cell migration [44].

Using high-throughput protein arrays combined with bioinformatics analysis, we have previously identified cortactin as a novel substrate of Pyk2 in invadopodia. Pyk2 binds cortactin via its PRR2 at invadopodia of invasive breast cancer cells, where it mediates EGF-induced phosphorylation both directly and indirectly via Src-mediated Arg activation, leading to free actin barbed end generation and subsequent actin polymerization in invadopodia, ECM degradation, and consequent tumor cell invasion [45]. Based on these findings, and on the synergism between *PTK2B* and *CTTN* genes in promoting breast cancer metastasis, we hypothesized that inhibition of the Pyk2/cortactin interaction could block invadopodia-mediated tumor cell invasiveness and consequent breast cancer metastasis. Here, we demonstrate that a cell-permeable peptide derived from the Pyk2-PRR2 blocks spontaneous lung metastasis in a xenograft mouse model by suppressing invadopodia-mediated invasiveness of breast tumor cells. The binding of the Pyk2-PRR2 peptide to the SH3 domain of cortactin was demonstrated in solution and the three-dimensional structure of this complex was determined using nuclear magnetic resonance (NMR). This native solution view of the SH3:PRR2 complex shows that cortactin interacts with Pyk2 via an extended binding interface, which includes a canonical SH3 interface and a second affinity-enhancing hydrophobic surface. Guided by this structure, we designed a mutant peptide unable to inhibit invadopodia maturation and consequent metastatic dissemination in mice, demonstrating the potential of our structure in facilitating the discovery of new Pyk2 inhibitors. Our findings offer a valuable molecular view of coratctin-Pyk2 interactions and suggest that their inhibition may be used as a novel strategy for blocking breast cancer metastasis.

## RESULTS

### Increased expression of Pyk2 and cortactin is associated with breast cancer metastasis and poor patient prognosis

We have previously shown that the Pyk2/cortactin interaction is critical for invadopodia-mediated breast tumor cell invasiveness [45]. We studied this further by integrating DNA, RNA, and protein expression data of breast cancer tissues from TCGA and stratifying them by hormone- and HER2-receptor status. A comparison of copy number alterations (CNA), mRNA expression, and protein expression demonstrated similar distribution among all hormone- and HER2-receptor statuses (Fig. 1A-1E), suggesting that Pyk2 and cortactin expression profile is not unique to any of the four primary molecular subtypes in breast cancer. To assess the involvement of Pyk2 and cortactin in metastatic dissemination, we analysed microarray expression levels of 1,650 breast cancer samples. The survival analysis showed significant association between poor distant metastasis free survival (DMFS) and high mRNA levels of *CTTN* (HR 1.56, 95% CI 1.29-1.93; *p*<0.001) but not with *PTK2B* (HR 1.06, 95% CI 0.86-1.30; *p*=0.62) (Fig. 1F-G). However, the combined effect of *PTK2B* and *CTTN* overexpression considerably reduced DMFS estimates (HR 1.72, 95% CI 1.29-2.30; *p*<0.001) with a synergy index (SI) of 1.13 (Fig. 1H), suggesting that Pyk2 and cortactin act synergistically to promote breast cancer metastasis, and that inhibition of their interaction may block this process.

**Fig. 1.**
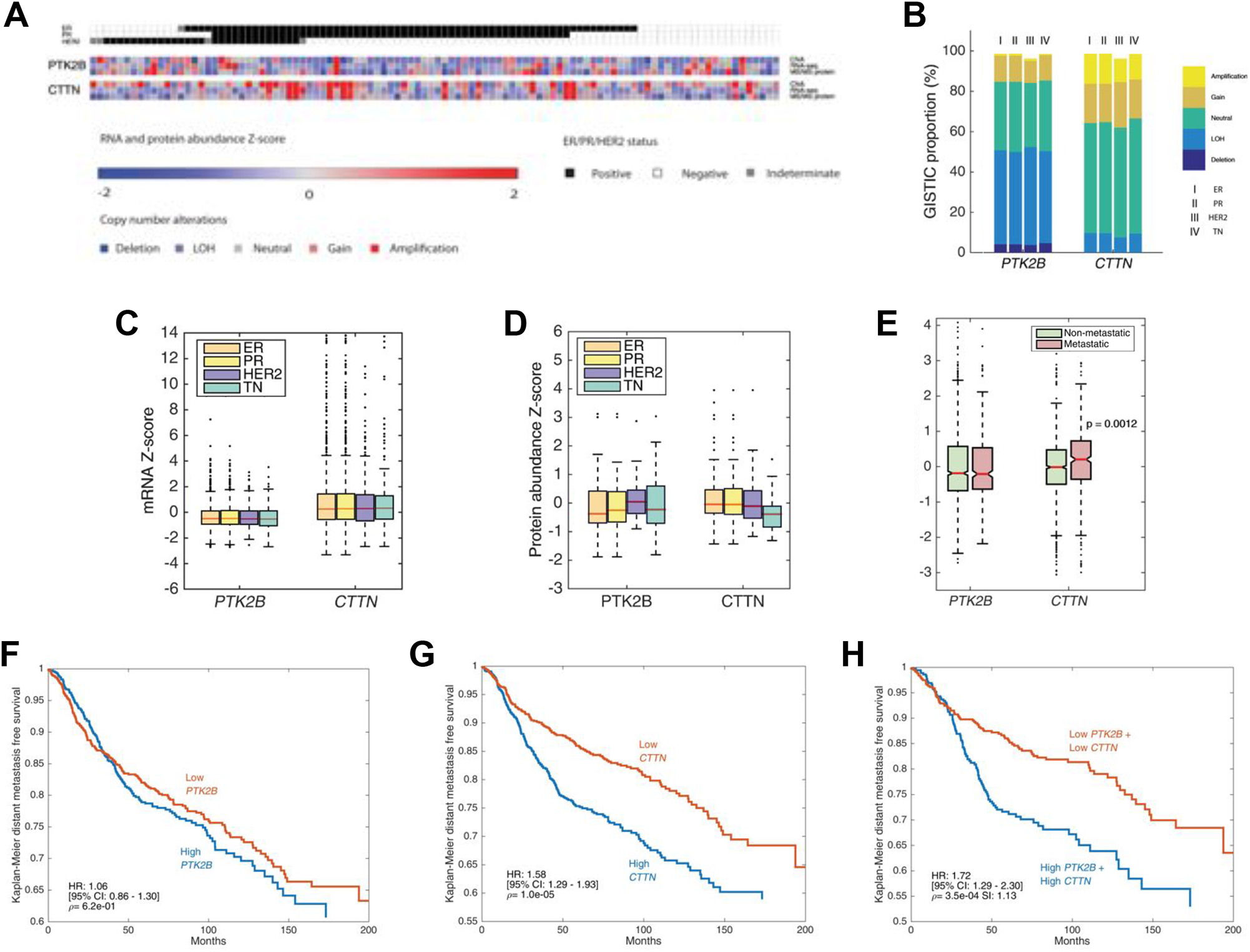
Increased expression of Pyk2 and cortactin is associated with breast cancer metastasis and poor patient prognosis. **(A)** Heat map of copy number alteration (CNA), mRNA expression, and protein expression of Pyk2 (PTK2B) and cortactin (CTTN) across 102 tumor samples based on the CPTAC database. ER, estrogen receptor positive; PR, progesterone receptor positive; HER2, HER2 positive; TN, triple negative. GISTIC, mRNA and protein abundance Z-scores are shown for each sample. **(B)** GISTIC analysis of 1,905 samples originated from TCGA with different hormone receptor statuses. **(C)** mRNA expression analysis of 1,905 samples originated from TCGA with different hormone receptor statuses. **(D)** Comparison of protein abundance in 102 samples from different hormone receptor statuses. **(E)** mRNA expression levels of *PTK2B* and *CTTN* in metastatic versus non-metastatic breast cancer patients is shown in boxplots. Metastasis was defined as occurrence of distant metastasis event within the follow-up period. The boxes in C, D and E are delimited by the lower and upper quartile, the horizontal red line indicates the sample median, the notches represent the 95% confidence interval (CI), and whiskers extend to the most extreme point which is no more than 1.5 times the interquartile range from the box. Outliers are shown as points beyond boxplot whiskers. P values were calculated using two-tailed Student’s *t*-test. **(F-H)** Kaplan-Meier curves of DMFS in 1,650 breast cancer cases. Microarray data (Affymatrix) were obtained from the NCBI Gene Expression Omnibus (GEO) data repository. Tumor samples were split into high and low expression groups based on mRNA gene expression with z-score cut-off value 0. Shown are DMFS survival curves for *PTK2B* (F), *CTTN* (G), and their combination (H). P values were calculated by log-rank test, hazard ratio (HR) and 95% CI are shown. Synergy index (SI) is indicated for evaluation of *PTK2B+CTTN* in (H).

### A peptide derived from the second proline-rich region of Pyk2 blocks spontaneous lung metastasis in an immune-competent xenograft mouse model

Given these results and considering our earlier findings that the Pyk2/cortactin-SH3 interaction is critical for invadopodia maturation and function [45], we hypothesized that its inhibition could block invadopodia-mediated breast cancer metastasis. Pyk2 peptides derived from the second (Pyk2-PRR2) or third (Pyk2-PRR3) proline-rich regions and a control scrambled peptide were synthesized and fused to an HIV-TAT sequence to increase their permeability into cells [46]. To examine their effect on breast cancer metastasis in mice, we used the highly tumorigenic 4T1 cell line that efficiently forms mammary tumors upon injection into the mammary fat pad of syngeneic immune-competent BALB/c mice and spontaneously metastasizes to multiple distant sites in a pattern similar to that of human breast cancer. Tumor-bearing mice were treated with the peptides by intraperitoneal injection for 8 days, followed by careful examination and quantification of lung metastases at the end of the treatment (Fig. 2A). As demonstrated in Fig. 2B-C, mice treated with Pyk2-PRR2 exhibited significantly fewer lung metastases than scrambled peptide- or Pyk2-PRR3-treated mice bearing equal-sized tumors.

**Fig. 2.**
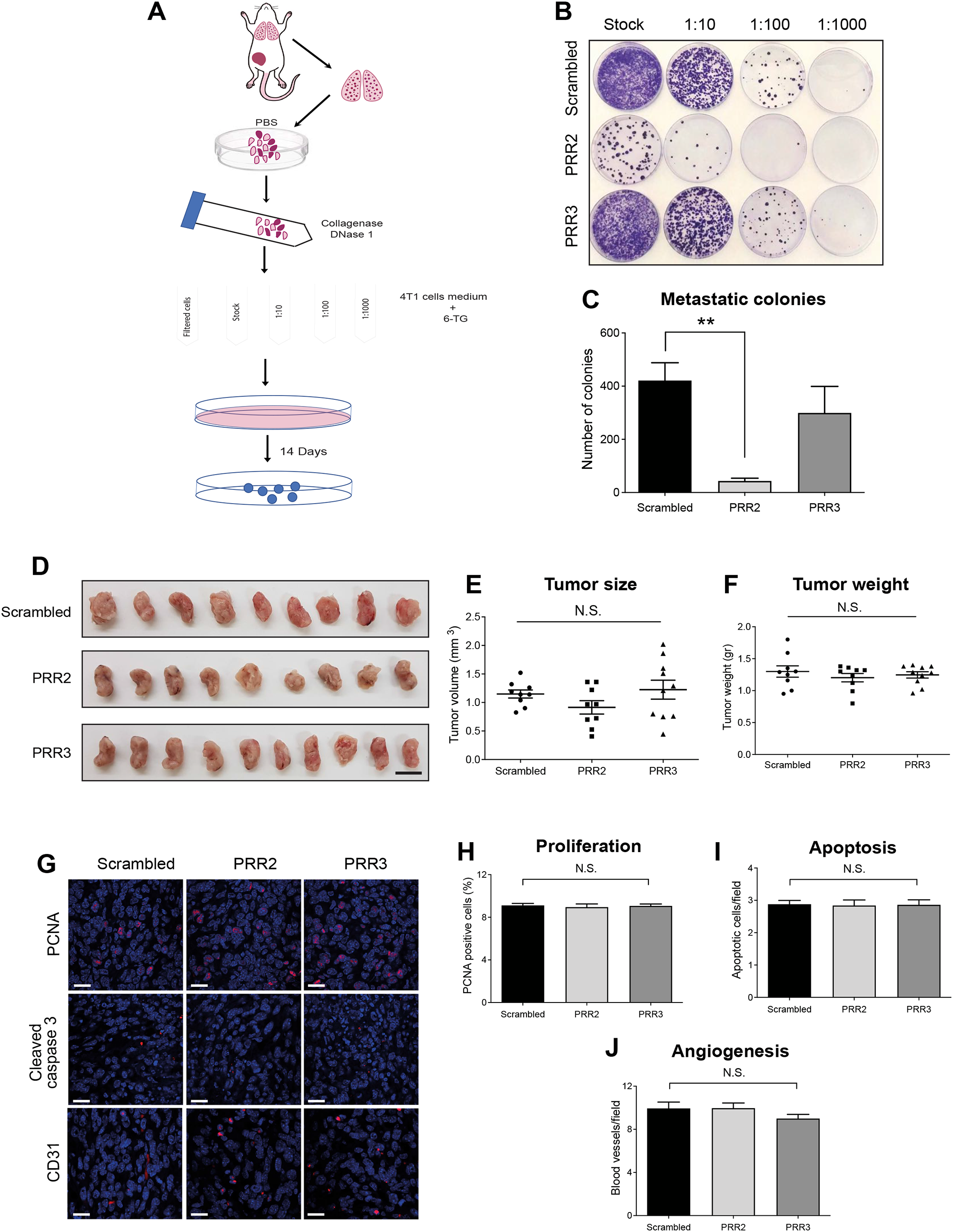
A cell-permeable peptide derived from the second proline-rich region of Pyk2 blocks spontaneous lung metastasis in an immune-competent xenograft mouse model. 4T1 cells were injected into the mammary fat pad of 8-week-old BALB/c female mice and allowed to grow for 12 days until the tumor reached the size of 100 mm^3^. On day 13 following injection, mice were treated by intraperitoneal injection with 10 mg/kg of scrambled control, Pyk2-PRR2, or Pyk2-PRR3 once a day for eight days. **(A)** Lungs were dissected at the end of the experiment, dissociated, and plated in different dilutions in a medium containing 6-thioguanine for 14 days. On day 14, 6-thioguanine colonies were stained with crystal violet, and colonies were counted. **(B)** Representative images of metastatic 4T1 resistant colonies from control, Pyk2-PRR2, and Pyk2-PRR3 treated mice. **(C)** Quantification of colonies from 1:10 dilution of dissociated lung cells. Each colony was originated from one metastatic cell. *n* = 7 (scrambled), *n* = 7 (Pyk2-PRR2), *n* = 8 (Pyk2-PRR3) mice per group. ** P < 0.01 by one-way ANOVA with Tukey’s post-hoc test. Error bars represent SEM. **(D-F)** Primary tumors were dissected at the end of the experiment (24 h following the last peptide injection), and tumor volume and weight were measured. *n* = 9 (scrambled), *n* = 9 (Pyk2-PRR2), *n* = 10 (Pyk2-PRR3) tumors per group. Scale bar, 1 cm. **(G)** Representative images of primary tumor sections stained with anti-PCNA (proliferation), anti-cleaved caspase 3 (apoptosis), and anti-CD31 (angiogenesis). Scale bar, 100 µm. **(H)** Quantification of PCNA positive cells (red/pink) normalized to DAPI positive cells (blue). The average percentage of proliferating cells in total cells per field is shown. **(I)** Quantification of apoptotic cells (cleaved caspase 3 positive cells). Shown is the number of apoptotic cells per field. **(J)** Quantification of CD-31 positive blood vessels. For all quantifications, *n* = 50 random fields from five tumors per condition.

To investigate these further, primary tumors were isolated and examined at the end of the treatment. As demonstrated in Fig. 2D-F, no significant differences in tumor size or weight were observed in mice treated with Pyk2-PRR peptides or the scrambled control. Accordingly, and as observed by immunofluorescent histological labeling, no significant differences were observed in proliferation, apoptosis, or angiogenesis in tumors from the different groups (Fig. 2G-J). Collectively, these data suggest that while none of the peptides suppressed primary tumor growth, the Pyk2-PRR2 peptide, but not Pyk2-PRR3 peptide, can suppress *in vivo* breast tumor metastasis in immune-competent mice bearing equal sized tumors.

### Pyk2-PRR2 suppresses MMP-dependent 3D invasiveness of breast tumor cells

Metastatic dissemination is initiated by the invasion of tumor cells through the tumor basement membrane and extracellular matrix (ECM). To study the effect of Pyk2 peptides on tumor cell invasiveness, we used the MDA-MB-231 cell line, a highly invasive, triple-negative basal-like human breast adenocarcinoma cell line that forms functional invadopodia in culture. Peptide permeability into MDA-MB-231 cells was validated using an immunofluorescence assay (Fig. S1A-B), showing all peptides penetrated the cells and were stable at least 24 h following incubation. An XTT assay on cells incubated with 0.1, 1, 5, and 10 μM concentrations of peptides showed that the three lower concentrations did not affect cell viability (Fig. S1C-F) and were therefore used in all further cellular studies. We examined the ability of the three peptides to inhibit the invasion of breast cancer cells embedded in a three-dimensional (3D) Matrigel matrix towards a scratch wound over 24 h. As demonstrated in Fig. 3D and Supplementary Movies 1-3, cells treated with the Pyk2-PRR2 peptide, but not with the Pyk2-PRR3 or control peptides, showed significantly reduced ability to close the gap by invading towards the wound. Significantly, ECM invasion was always fully inhibited by MMP inhibitor GM6001 (Fig. 3H-J), confirming that this invasion is MMP-mediated and that the Pyk2-PRR2 peptide inhibits MMP-dependent tumor cell invasiveness.

**Fig. 3.**
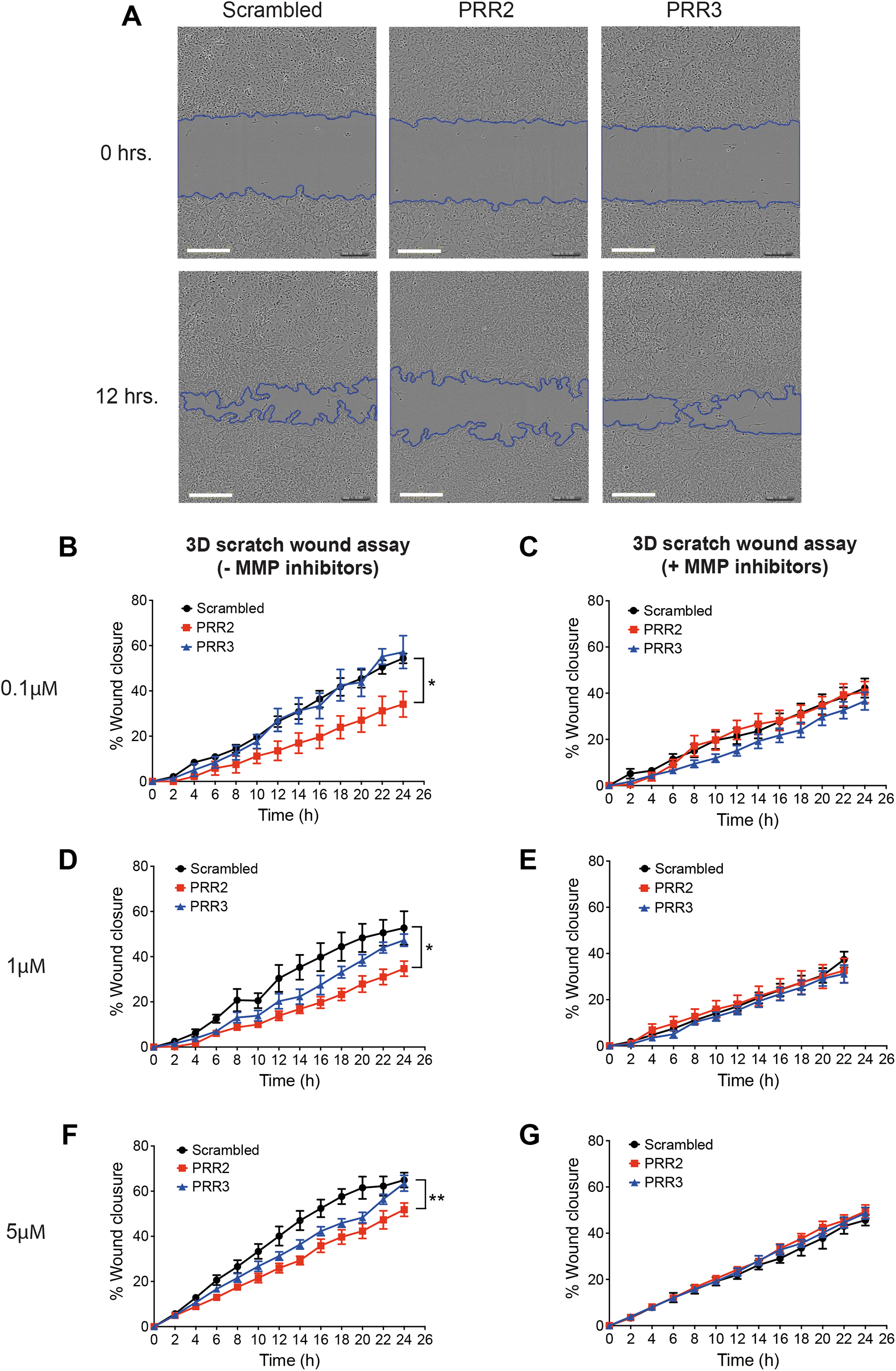
Pyk2-PRR2 suppresses MMP-dependent 3D invasiveness of breast tumor cells. **(A)** Representative images from 3D scratch-wound assay movies of MDA-MB-231 cells treated with Pyk2-PRR or scrambled peptides at time 0 (upper panels) and 12 h (lower panels). Scale bar, 300 µm. **(B, D, F)** Quantification of 3D directed motility towards a wound in Matrigel containing embedded MDA-MB-231 cells treated with Pyk2-PRR2 (red), Pyk2-PRR3 (blue), or scrambled control (black) peptides. **(C, E, G)** Quantification of 3D directed motility towards a wound in Matrigel containing embedded MDA-MB-231 cells treated with Pyk2-PRR2 (red), Pyk2-PRR3 (blue), or scrambled control (black) peptides and normalized to MMP inhibitor-treated values. *n* = 3 independent experiments, each experiment was performed in triplicates. * P < 0.05 (between scrambled and PRR2 and scrambled and PRR3 in B; between scrambled and PRR2 in D), ** P < 0.01 (between scrambled and PRR2 in F), as determined by two-way ANOVA followed by Bonferroni post-hoc test. Error bars represent SEM.

Further evidence of the Pyk2-PRR2 effect upon tumor cell invasiveness was provided by a 2D random migration assay. Invasiveness is controlled by coordinating MMP-dependent invadopodia-mediated invasion and focal adhesion-mediated motility [6, 47], and we have previously shown that Pyk2 localizes to these regions and that knockdown of Pyk2 inhibits both MMP-dependent invasion and two-dimensional (2D) motility [45]. Following accumulated and Euclidean distances as well as velocity over 12 h, cells treated with Pyk2-PRR2 at all concentrations and with Pyk2-PRR3 at the highest concentration showed reduced 2D motility over an ECM substrate compared to the control peptide (Fig. S2). Thus, Pyk2-PRR2, and to a lesser extent Pyk2-PRR3, regulate 2D motility and inhibit tumor cell invasiveness.

### The Pyk2-PRR2 peptide inhibits the interaction between Pyk2 and cortactin in invadopodia and consequent invadopodium maturation and function in breast tumor cells

Breast cancer metastasis has previously been correlated *in vivo* with invadopodia maturation and function [7, 8], and in mouse models and human patients with invadopodia assembly and function [5-7, 48]. Since we have shown that Pyk2 directly binds cortactin in invadopodia of breast cancer cells via its PRR2 [45], and considering the above findings, we hypothesized that the Pyk2-PRR2 peptide binds the cortactin SH3 domain, thus inhibiting the Pyk2-cortactin interaction in invadopodia. Binding of Pyk2-PRR peptides to purified cortactin-SH3 was determined by following ligand-induced chemical shift changes along the course of a titration isotherm. Pyk2-PRR2 and Pyk2-PRR3 ligands exhibited affinities of *K*_D_ = 40 ± 5 and *K*_D_ = 650 ± 10 μM, respectively, while a scrambled control peptide did not display significant affinity to the SH3 domain (*K*_D_ > 1 mM) (Fig. 4A). Largest Pyk2-PRR2-induced changes were observed for residues I521, A539 and the W525 indole cross-peak, consistent with a canonical binding mode. Thus, the Pyk2-PRR2 peptide binds to cortactin SH3 *in vitro* with a significantly (16-fold) higher affinity than Pyk2-PRR3. We also measured the ability of Pyk2-PRR2 to block the access of Pyk2 to cortactin in invadopodia in Förster resonance energy transfer (FRET) acceptor photobleaching experiments. These demonstrated that the Pyk2-PRR2, but not Pyk2-PRR3 or a scrambled peptide, significantly inhibited the interaction between Pyk2 and cortactin in invadopodia (Fig. 4B-C).

**Fig. 4.**
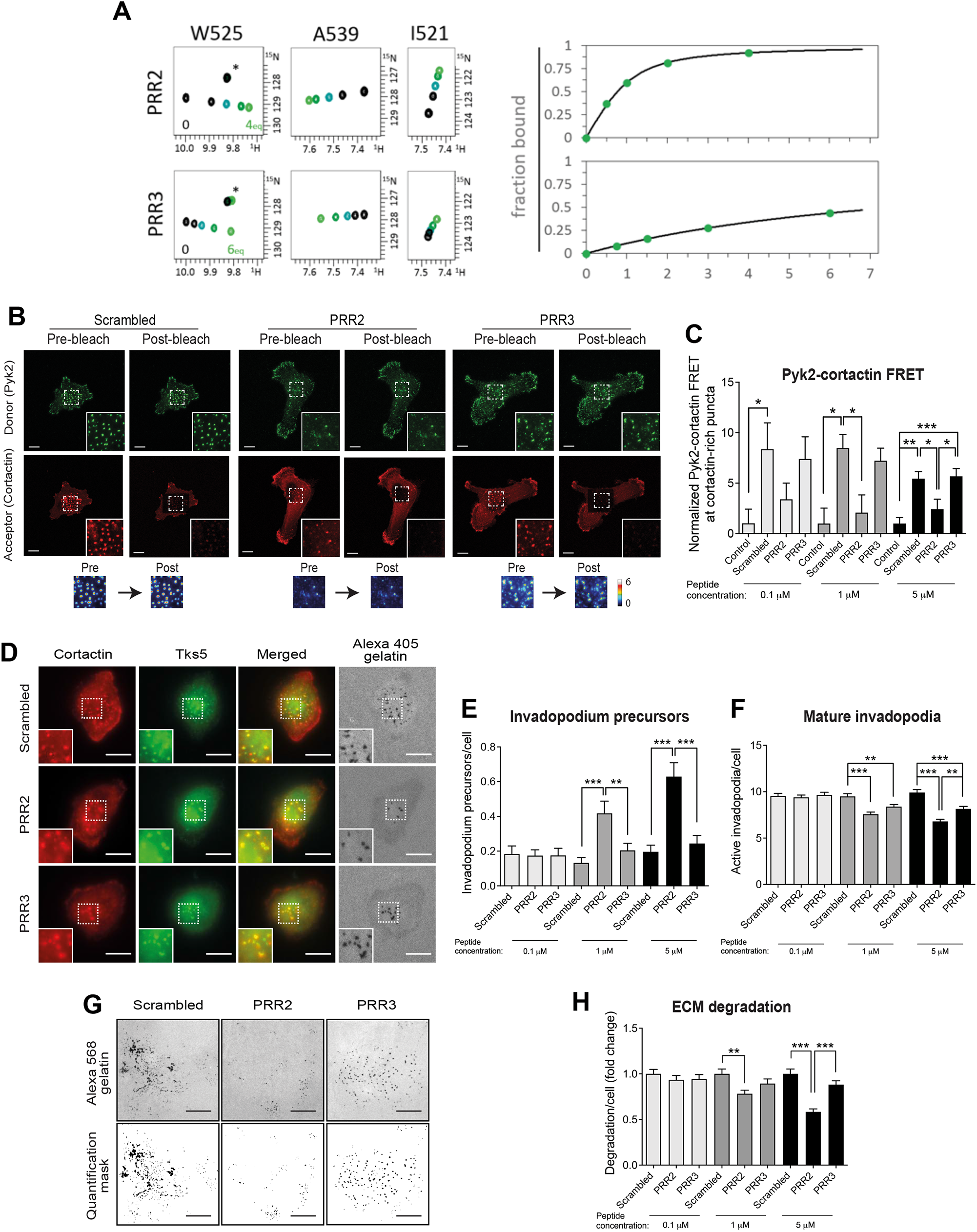
Pyk2-PRR2 peptide inhibits Pyk2-cortactin interactions in invadopodia and consequent invadopodium maturation and function in breast tumor cells. **(A)** Left, NMR determination of SH3 binding of Pyk2 peptides PRR2 and PRR3. Details from the SH3 ^1^H,^15^N-HSQC spectra acquired along a titration isotherm with focus on residues W525 (indole sidechain) and I521, A539 (backbone amide). Spectra were acquired at 0:1 (black), 0.5:1, 1:1, 2:1 and 4:1 (light green) mol:mol PRR2:SH3 and 0:1 (black), 0.75:1, 1.5:1, 3:1 and 6:1 (light green) mol:mol PRR3:SH3. In the W525 panel, the asterisk indicates a different (W526 indole) unaffected cross-peak. Right, binding isotherms showing the fraction of bound SH3 as a function of the PRR:SH3 mol:mol ratio, and the fitted curve providing the dissociation constant. Not shown is scrambled PRR2, for which a 6:1 peptide:SH3 mol:mol ratio exhibited a small effect on peak position, projecting to a low (*K*_D_ > 1 mM) affinity for this peptide. **(B)** Representative images of Pyk2-cortactin FRET in invadopodia of MDA-MB-231 cells stably expressing cortactin-TagRFP, which were treated with scrambled peptide, Pyk2-PRR2, or Pyk2-PRR3 and labeled for pY402-Pyk2 (green) and cortactin (red). Scale bar, 10 µm. **(C)** Quantification of FRET between Pyk2-pY402 and cortactin in mature, matrix-degrading invadopodia of MDA-MB-231 cells treated with different peptide concentrations. *n* = 42-66 invadopodia per group from three different experiments. **(D)** MDA-MB-231 cells were plated on Alexa 405-labeled gelatin and incubated with different concentrations of Pyk2-PRR or control scrambled peptides for 4 h, fixed, and labeled for cortactin (red) and Tks5 (green). Boxed regions and insets depict co-localization of cortactin and Tks5 as markers of invadopodium precursors and Alexa 405 gelatin as a marker of mature invadopodia. Scale bar, 10 µm. **(E)** Quantification of invadopodium precursors defined by co-localization of cortactin and Tks5. *n* = 136-153 (scrambled), *n* = 143-155 (Pyk2-PRR2), *n* = 156-160 (Pyk2-PRR3) cells from three different experiments. **(F)** Quantification of mature (active) invadopodia, defined by co-localization of cortactin and Tks5 with degradation regions. *n* = 138-153 (scrambled), *n* = 143-155 (Pyk2-PRR2), *n* = 156-160 (Pyk2-PRR3) cells from three different experiments. **(G)**MDA-MB-231 cells were plated on Alexa Fluor 568 FN/gelatin matrix in the presence of Pyk2-PRR or control peptides and allowed to degrade for 24 h. Shown are representative images (upper panels) and quantification masks (lower panels) of degradation areas. Scale bar, 25 µm. **(H)**Quantification of matrix degradation by peptide-treated cells. *n* = 91-127 (scrambled), *n* = 91-123 (Pyk2-PRR2), *n* = 92-105 (Pyk2-PRR3) fields per group from three independent experiments. *P < 0.05, ** P < 0.01, *** P < 0.001, as determined by one-way ANOVA followed by Tukey’s post-hoc test. Error bars represent SEM.

To examine whether Pyk2-PRR2 regulates the initial assembly of invadopodium precursors, MDA-MB-231 cells were treated with Pyk2-PRR2, Pyk2-PRR3, or scrambled control peptide, plated on fluorescently labeled gelatin matrix and labeled for Tks5 and cortactin as invadopodium precursor markers (Fig. 4D). The significant increase in immature invadopodium precursors, along with a significant decrease in mature, matrix-degrading invadopodia observed in Pyk2-PRR2-treated cells implied that this peptide blocks the maturation and activation of pre-assembled invadopodium precursors in breast tumor cells (Fig. 4E, F). The minor effect of Pyk2-PRR3 at higher peptide concentrations in both 2D motility and in invadopodia formation and maturation is consistent with its weaker binding affinity to cortactin-SH3 and lower similarity to the cortactin consensus motif.

At the final maturation stage invadopodia locally degrade the ECM by secretion and activation of MMPs [19, 22], allowing invasive cancer cells to leave the primary tumor and penetrate blood vessels. Breast tumor cells were plated on fluorescent fibronectin/gelatin matrix and allowed to degrade the matrix for 24 h in the presence of increasing concentrations of control, Pyk2-PRR2, or Pyk2-PRR3 peptides (Fig. 4G). Quantification of the degradation area revealed a significant decrease in the ability of Pyk2-PRR2 treated cells to degrade the matrix in all concentrations tested (Fig. 4H). Taken together, these data suggest that the Pyk2-PRR2 peptide inhibits invadopodia maturation and activation and consequent MMP-dependent matrix degradation, which further supports our *in vitro* 3D invasion and our *in vivo* metastasis phenotypes.

### Pyk2-PRR2 inhibits cortactin tyrosine phosphorylation and consequent barbed end generation in invadopodia of breast cancer cells

Tyrosine phosphorylation of invadopodial cortactin mediates invadopodia maturation and activation, and involves the Pyk2 kinase, both directly and indirectly via Src-mediated Arg activation [45]. To see whether Pyk2-PRR peptides can inhibit this process in breast cancer cells, MDA-MB-231 cells stably expressing Tag-RFP-cortactin were treated with Pyk2-PRR or control peptides and further induced to form invadopodia by EGF. Cells were then fixed and labeled with phosphorylation-specific antibodies for cortactin tyrosine Y421. As demonstrated in Fig. 5A-B, phosphorylation was completely abolished in Pyk2-PRR2-treated cells at all concentrations, suggesting that Pyk2-PRR2 regulates the maturation and functional activation of invadopodia by inhibiting cortactin tyrosine phosphorylation. It is likely that this occurs by interrupting the binding of tyrosine kinases that normally phosphorylate cortactin, such as Pyk2 and Arg, to the SH3 domain of cortactin. To confirm these results, we also examined the effect of Pyk2-PRR2 on generation of free actin barbed ends, a known consequence of phosphorylation required for actin polymerization followed by pushing of the invadopodium membrane into the ECM and penetration of tumor cells through it [22]. We stimulated peptide-treated cells with EGF to synchronize the generation of free actin barbed ends within invadopodia, then quantified the generation of free actin barbed ends specifically within these structures (Fig. 5C). Whereas EGF stimulation generated a 1.3-1.8-fold increase in barbed end intensity in control- and Pyk2-PRR3-treated tumor cells, treatment with the Pyk2-PRR2 peptide completely disrupted generation of free actin barbed ends at invadopodia at all concentrations (Fig. 5D). Thus, inhibition of phosphorylation appears to be involved in the mechanism of Pyk2-PRR2 peptide blocking of invadopodial maturation in breast tumor cells.

**Fig. 5.**
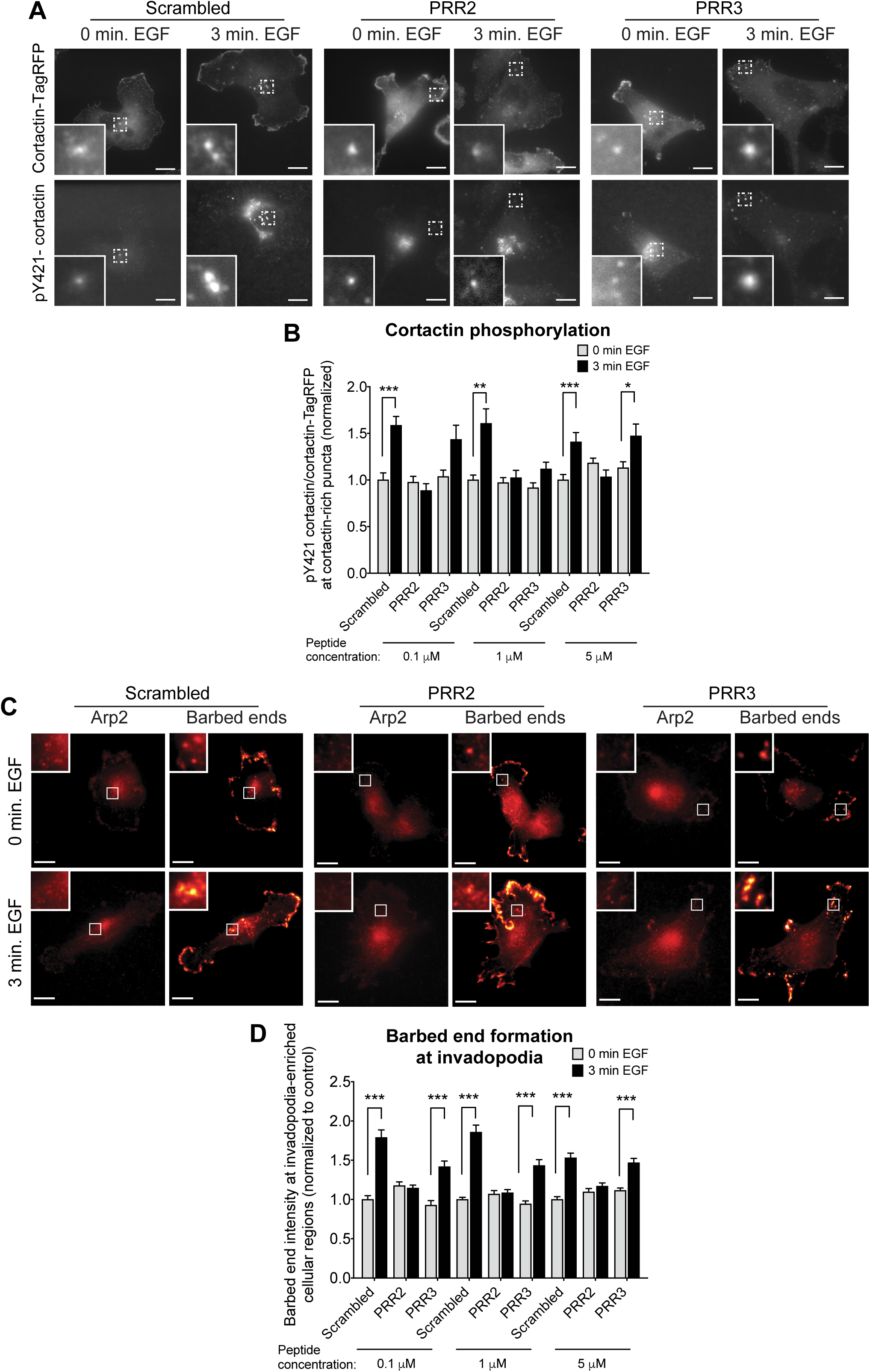
Pyk2-PRR2 inhibits cortactin tyrosine phosphorylation and consequent barbed end formation in invadopodia. **(A)** MDA-MB-231 cells stably expressing cortactin-TagRFP (upper panels) were treated with Pyk2-PRR or control scrambled peptides, plated on FN/gelatin matrix, starved, and stimulated with EGF. Cells were then fixed and labeled for phosphorylated cortactin (anti-pY421-cortactin; lower panels) before (0 min.) or after (3 min.) EGF stimulation. Scale bar, 10 µm. **(B)** Quantification of pY-421-cortactin/cortactin-TagRFP signal at cortactin-rich puncta. n = 42-129 (scrambled -EGF), *n* = 43-131 (scrambled +EGF), *n* = 42-142 (Pyk2-PRR2 -EGF), *n* = 42-116 (Pyk2-PRR2 +EGF), *n* = 41-92 (Pyk2-PRR3 -EGF), *n* = 41-100 (Pyk2-PRR3 +EGF) puncta per group from three independent experiments. **(C)** MDA-MB-231 cells were treated with Pyk2-PRR or control scrambled peptides, plated on FN/gelatin matrix, starved, and left untreated (0 min EGF) or stimulated with EGF for three min (3 min EGF). Cells were fixed and labeled for Arp2 (left panels) as invadopodia marker and biotin-actin (right panels) as a marker for newly formed barbed ends. Scale bar, 10 µm. **(D)** Quantification of free actin barbed ends as measured by the average biotin-actin intensity at stimulated invadopodia containing Arp2. *n* = 107-246 invadopodia per group from three different experiments. * P < 0.05, ** P < 0.01, *** P < 0.001, as determined by two-way ANOVA followed by Bonferroni post-hoc test. Error bars represent SEM.

### The structure of the Pyk2-PRR2/cortactin-SH3 complex reveals an extended binding interface

With extensive evidence of the Pyk2-PRR2/cortactin-SH3 interaction and its potent role in regulating invadopodia-mediated metastasis, we were motivated to characterize the complex on the molecular level to identify critical residues which contribute to its affinity. The NMR-based SH3/PRR2 complex structure was determined using distance constraints established through homonuclear and filtered-edited heteronuclear NOESY experiments and dihedral angle constraints derived from analysis of chemical shifts (Fig. 6; see also Supplementary Table 1). The SH3 domain adopts the well-known double β-sheet fold with the long β2-strand stitching the two sheets and forming the SH3 hydrophobic core. However, the complex interface revealed a critical extension of the canonical binding surface, which has not been previously reported for cortactin complexes. While the canonical interface is formed by seven SH3 residues (Y497, Y499, E506, W525, P538, N540, Y541) which bind to the hallmark class II polyproline helix ^207^PPPKPSR^213^, an additional interaction surface is formed between SH3 residues I521, D522, W525 and L536, and the Pyk2 extension ^214^PKY^216^. In the canonical interface, the dipeptide motifs ^207^PP^208^ and ^210^KP^211^ are inserted deepest in the two canonical SH3 binding grooves formed by residues Y497-Y541 and Y499-W525-P538-Y541, respectively. Concomitantly, residues P^211^SRP^214^ form a second ‘arch’ extending over the W525 indole ring, and residues ^214^PKY^216^ contact the second binding surface (surface-II), including I521 and L536 (hydrophobic interaction with Y^216^), W525 (interaction with P^214^), and D522 (electrostatic interaction with K^215^). These two SH3-PRR2 contacts contribute 844 and 424 Å^2^, respectively, to the overall binding surface (1,268 Å^2^) (Fig. 6A-B).

**Fig. 6.**
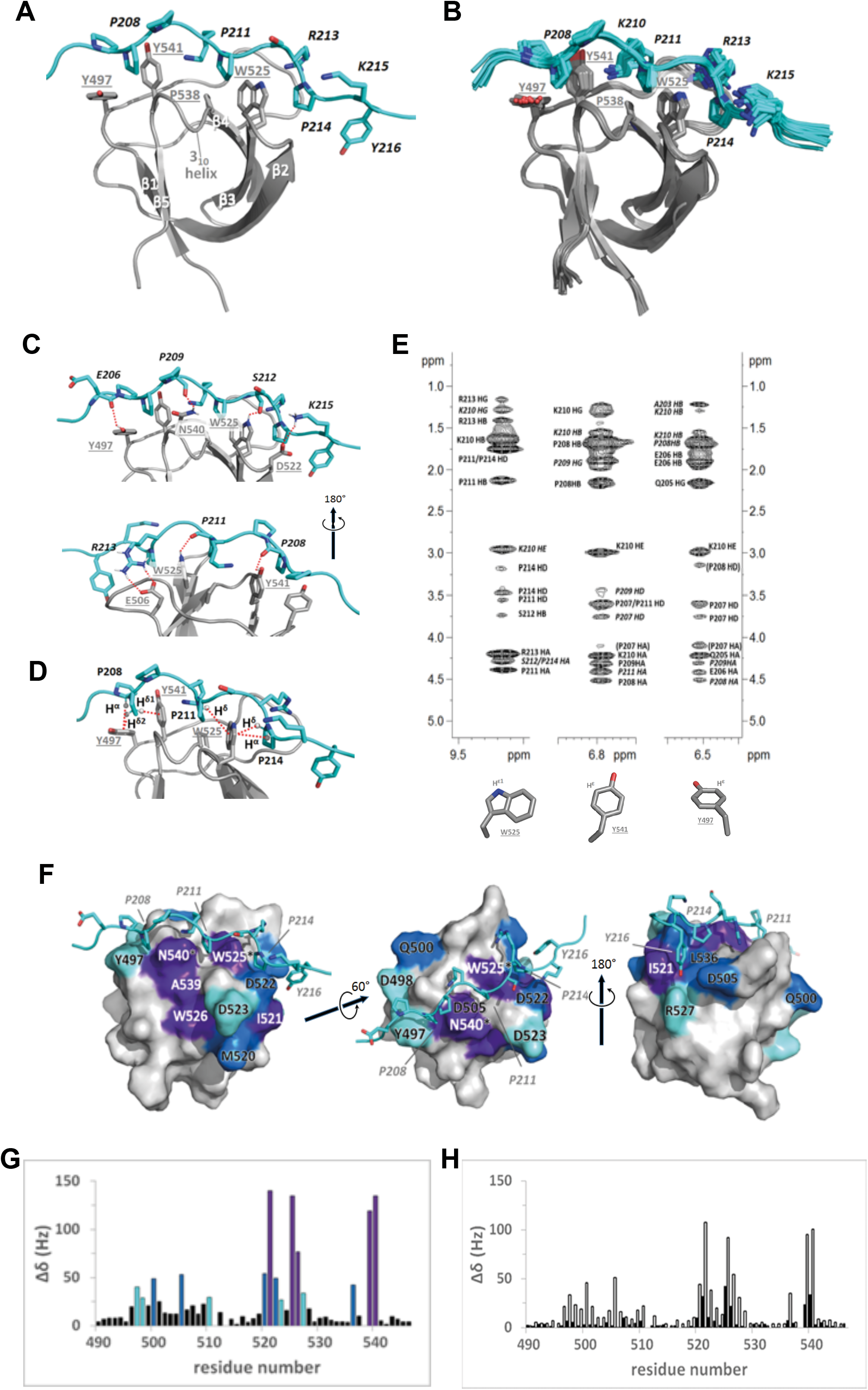
Structure of the cortactin-SH3:Pyk2-PRR2 complex. **(A)** Ribbon-representation of the complex showing the SH3 domain (grey, residue numbers in underlined grey) and the PRR2 peptide (cyan, residue numbers in black and *italics*). (**B)** Overlay of 20 low-energy structures of the complex with the SH3 domain aligned, exhibiting an excellent structure definition. Coloring and annotations as in A. (**C)** Network of hydrogen bonds (red dotted lines) contributing to the SH3:PRR2 affinity. Indicated hydrogen bonds were present in over 70% of structures as determined by analysis using VADAR 1.8. For clarity, the complex is shown from the front (top) and back (bottom). (**D)** Focus on possible CH-pi interactions between aromatic pi-systems and polar CH bonds (α- and δ-protons of prolines). (**E**) NOESY cross-peaks emanating from the SH3:PRR2 interface, involving W525 H^ε1^ (left), Y541 H^ε^ (center), and Y497 H^ε^ (right). Annotations in italics indicate cross-peaks for which distance constraints were extended due to side-chain flexibility. Annotations in parentheses indicate cross-peaks excluded from the analysis due to ambiguous assignment. **(F)** Three views of the complex showing the interaction of PRR2 (cyan ribbon) with the SH3 surface. SH3 residues are colored according to their PRR2-induced chemical shift perturbation (CSP); the third, second, and top deciles (each spanning five residues) are colored in cyan, blue, and purple, respectively. Asterisks denote changes in the side-chain cross-peaks of Trp and Asn residues. **(G)** Summary of PRR2-induced CSPs along the SH3 sequence, color-coding as in panel A, values for W525 and N540 are for side-chain CSPs. (**H)** PRR2-induced CSPs for the Y216A mutant (light grey bars) and the Δ214-220 truncated peptide (dark grey bars). Changes are far more pronounced for the truncation mutant, supporting the importance and interactions of surface II.

Three main features contribute to the affinity of this complex. First, a network of hydrogen bonds stabilizes the complex, including the P^208^ CO with the Y541 sidechain hydroxyl, the P^209^ CO with the N540 polar sidechain, and P^211^/S^212^ CO groups with the W525 indole NH. In addition, stabilizing electrostatic contacts are found between R^213^ and the acidic specificity pocket of SH3 residues D505/E506 and between K^215^ and D522 in surface II (Fig. 6C). Second, the P^208^/P^211^/P^214^pyrrolidine rings adopt an orientation parallel to the SH3 groove-lining aromatic residues W525 and Y541, thus favoring C-H-π interactions between the electron-rich aromatic rings and the partial positive charge of polarized Cα-Hα and Cδ-Hδ bonds in prolines (Fig. 6D) [49]. Finally, close packing of hydrophobic surfaces is found for the three inserted proline residues and residue Y216 with the methyl groups of I521 and L536.

The two-surface interaction between SH3 and PRR2 was further confirmed in the NMR chemical shift perturbation (CSP) data, as the strongest effects were observed for both the canonical binding surface (Y497, Q500, D505, W525, A539, N540) as well as residues near the second surface (M520, I521, D522, W526) (Fig. 6F and S3,4). Furthermore, the Y216A mutation reduced affinity 3.5-fold, and the Δ214-220 deletion mutant demonstrated a 27-fold loss of affinity and most significant CSP reductions at surface II residues I521 and L536 (Fig. 6G-H). Overall, we found the cortactin-Pyk2 interaction to span an extended proline-rich motif, including the canonical class II binding sequence and an additional C-terminal tripeptide extension with a second contributing binding surface.

### A super-mutant peptide cannot inhibit the Pyk2-cortactin interaction at invadopodia and spontaneous lung metastasis in a xenograft mouse model

We synthesized a super-mutant Pyk2-PRR2 peptide, in which five critical amino acids were mutated (P208/211/214A, R213Q, K215Q). The loss of SH3 affinity of this peptide was verified by NMR, in which no detectable affinity could be observed (estimated *K*_D_ > 1 mM). Similarly, FRET acceptor photobleaching experiments demonstrated that, unlike the Pyk2-PRR2 peptide, this super-mutant peptide does not significantly inhibit the Pyk2-cortactin interaction in invadopodia of breast tumor cells at any of the concentrations tested (Fig. 7A, B). We then repeated the invadopodia assembly assay on fluorescently labeled gelatin matrix for control, Pyk2-PRR2, and super-mutant peptide, and labeled for Tks5 and cortactin (Fig. 7C). While no significant effect was observed for the super-mutant peptide in the number of invadopodium precursors formed in cells, a small but significant reduction was observed in the number of mature degrading invadopodia (Fig. 7D, E), and consequently, in the levels of ECM degradation (Fig. 7F, G, compared to control peptide in both cases). Thus, although the super-mutant peptide cannot bind to the cortactin-SH3 domain *in vitro* or inhibit the cortactin-Pyk2 interaction in invadopodia, it can slightly (albeit less than Pyk2-PRR2) affect invadopodia maturation and subsequent ECM degradation. This residual effect may result from non-specific low-affinity binding of the remaining proline and tyrosine residues to the additional hydrophobic groove within the SH3 domain of cortactin.

**Fig. 7.**
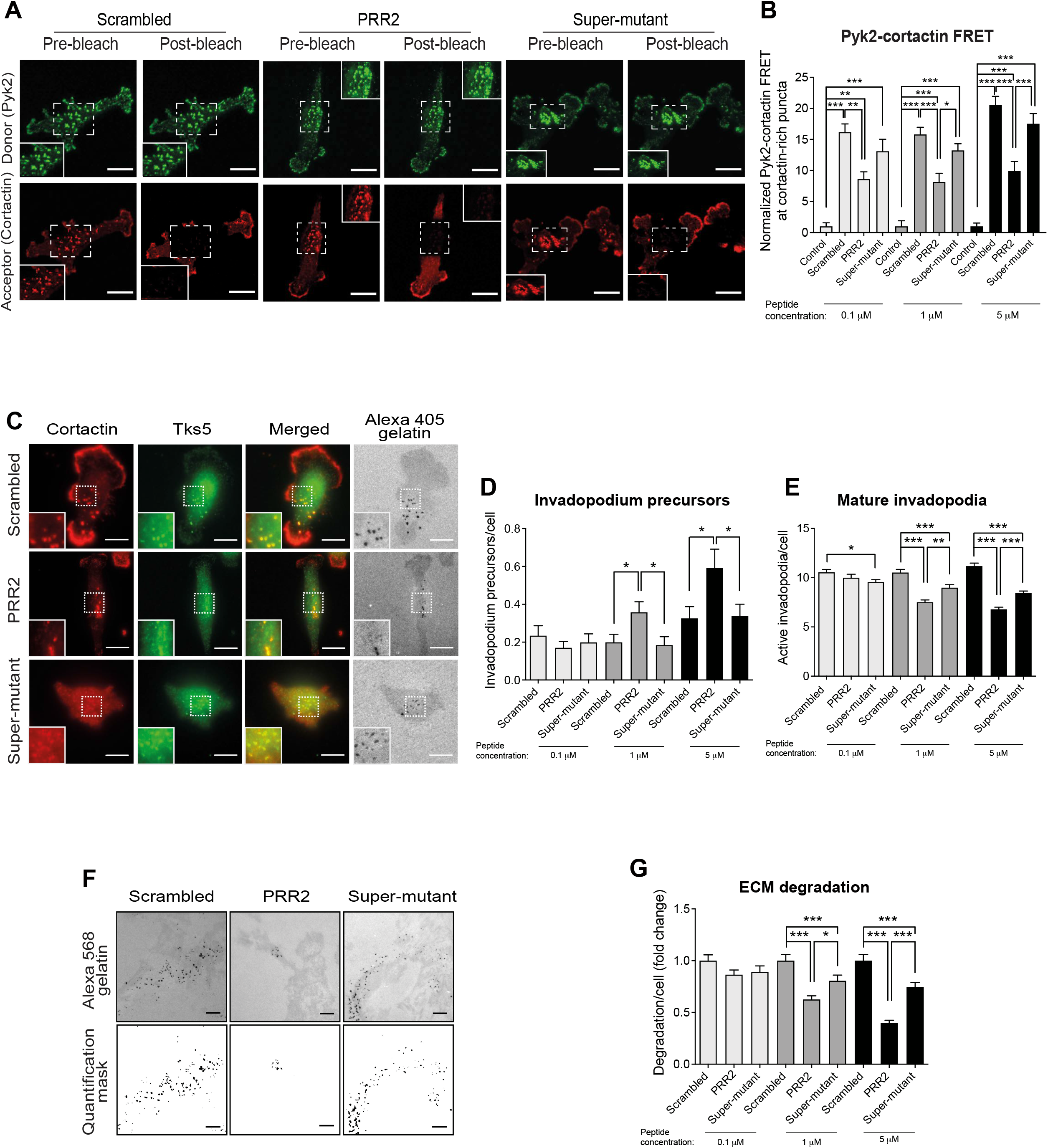
A Pyk2-PRR2 super-mutant peptide does not inhibit the interaction between Pyk2 and cortactin at invadopodia. **(A)** Representative images of Pyk2-cortactin FRET in invadopodia of MDA-MB-231 cells stably expressing cortactin-TagRFP-WT and treated with either scrambled peptide, Pyk2-PRR2, or Pyk2 super-mutant peptides and labeled for pY402-Pyk2 (green) and cortactin (red). **(B)** Quantification of FRET between Pyk2-pY402 and cortactin in mature, matrix-degrading invadopodia of MDA-MB-231 cells treated with different peptide concentrations. *n* = 68-72 invadopodia per group from three different experiments. **(C)** MDA-MB-231 cells were plated on Alexa 405-labeled gelatin and incubated with different concentrations of control, Pyk2-PRR2, or Pyk2 super-mutant for 4 h, fixed, and labeled for cortactin (red) and Tks5 (green). Boxed regions and insets depict co-localization of cortactin and Tks5 as markers of invadopodium precursors and Alexa 405 gelatin as a marker of mature invadopodia. Scale bars: 10 µm (main images), 2 µm (insets). **(D)** Quantification of invadopodium precursors defined by co-localization of cortactin and Tks5. *n* = 158-178 (scrambled), *n* = 141-154 (Pyk2-PRR2), *n* = 141-174 (Pyk2 super-mutant) cells from three independent experiments. **(E)** Quantification of mature (active) invadopodia, defined by co-localization of cortactin and Tks5 with degradation regions. *n* = 158-178 (scrambled), *n* = 141-154 (Pyk2-PRR2), *n* = 141-174 (Pyk2 super-mutant) cells from three independent experiments.**(F)** MDA-MB-231 cells were plated on Alexa Fluor 568 FN/gelatin matrix in the presence of control scrambled, Pyk2-PRR2, or Pyk2 super-mutant and allowed to degrade for 24 h. Shown are representative images (upper panels) and quantification masks (lower panels) of degradation areas. Scale bar, 25 µm. **(G)** Quantification of matrix degradation by peptide-treated cells. *n* = 115 (scrambled), *n* = 106 (Pyk2-PRR2), *n* = 120 (Pyk2 super-mutant) cells from three independent experiments. ** P < 0.01, *** P < 0.001, as determined by one-way ANOVA followed by Tukey’s post-hoc test. Error bars represent SEM.

We also repeated the xenograft model assay for the Pyk2 super-mutant peptide, treating tumor-bearing mice with a cell-permeable Pyk2 super-mutant (with a scrambled peptide as control) by intraperitoneal injection for eight days. As above, no significant differences were detected in primary tumor growth in all groups (Fig. 8A-C), which was supported by similar proliferation, apoptosis, and angiogenesis traits of tumors in all peptide-treated mice (Fig. 8D-G). However, while Pyk2-PRR2 significantly inhibited lung metastasis, no significant differences in colony number, indicating lung metastasis, were observed between scrambled control and the Pyk2 super-mutant-treated tumor-bearing mice (Fig. 8H-I). These data demonstrate that, while the Pyk2 super-mutant peptide may have a residual effect on invadopodia maturation and function *in vitro*, it fails to inhibit spontaneous lung metastasis of breast tumor cells *in vivo*.

**Fig. 8.**
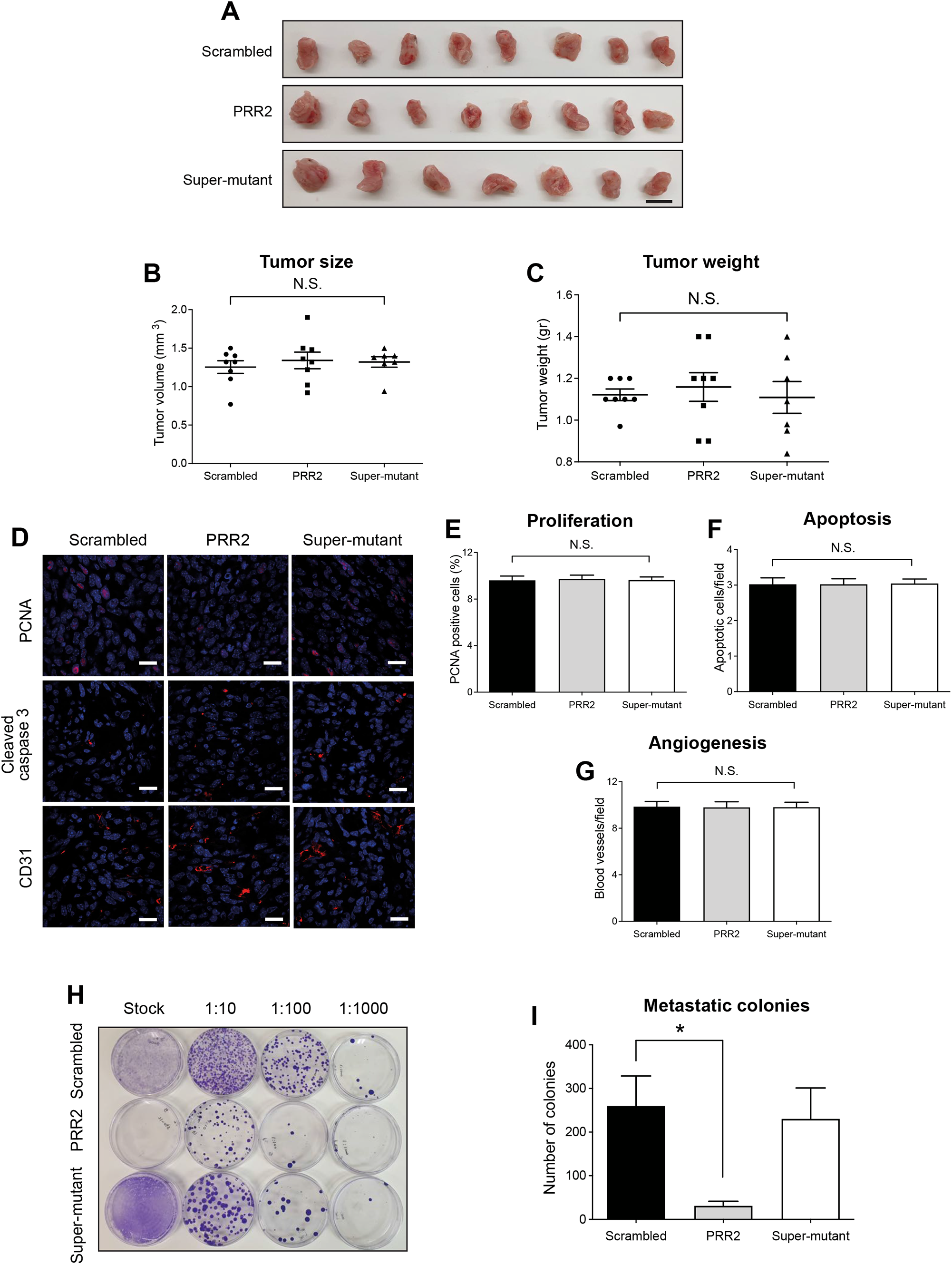
The Pyk2 super-mutant peptide does not block spontaneous lung metastasis in a xenograft mouse model. 4T1 cells were injected into the mammary fat pad of 8-week-old BALB/c female mice and allowed to grow for 12 days until the tumors reached an average size of 100 mm^3^. On day 13 following injection, mice were treated by intraperitoneal injection with 10 mg/kg of scrambled control, Pyk2-PRR2, or Pyk2 super-mutant once a day for eight days. **(A-C)** Primary tumors were dissected at the end of the experiment (24 h following the last peptide injection), and tumor volume and weight were measured. *n* = 8 (scrambled), *n* = 8 (Pyk2-PRR2), *n* = 7 (Pyk2 super-mutant) tumors per group. Scale bar, 1 cm. **(D)** Representative images of primary tumor sections stained with anti-PCNA (proliferation), anti-cleaved caspase 3 (apoptosis), and anti-CD31 (angiogenesis). Scale bar, 100 µm. **(E)** Quantification of PCNA positive cells (red/pink) normalized to DAPI positive cells (blue). The average percentage of proliferating cells in total cells per field is shown. **(F)** Quantification of apoptotic cells (cleaved caspase 3 positive cells). Shown is the number of apoptotic cells per field. **(G)** Quantification of CD-31 positive blood vessels. *n* = 44 (scrambled), *n* = 47 (Pyk2-PRR2), *n* = 47 (Pyk2 super-mutant) random fields from 5 tumors per condition. **(H)** Representative images of metastatic 4T1 resistant colonies from control, Pyk2-PRR2, and Pyk2 super-mutant treated mice. **(I)** Quantification of colonies from 1:10 dilution of dissociated lung cells. Each colony was originated from one lung metastatic cell. *n* = 8 (scrambled), *n* = 8 (Pyk2-PRR2), *n* = 6 (Pyk2 super-mutant) mice per group. * P < 0.05 by one-way ANOVA followed by Tukey’s post-hoc test. Error bars represent SEM.

## DISCUSSION

An estimated 90% of deaths from breast cancer are a consequence of metastatic disease. While treatments that slow tumor growth are commonly in use, no treatment that blocks metastatic breast cancer currently exists. Here we describe a Pyk2-derived cell-permeable inhibitor of invadopodia maturation in breast tumor cells and consequent metastasis in an immune-competent mouse model. Our first molecular view of the cortactin SH3:PRR2 complex demonstrates that they interact via two major interfaces, suggests amino acids that are critical for Pyk2-PRR2 peptide binding, and forms the basis for design of more potent peptide inhibitors for blocking breast cancer metastasis. Although two earlier crystal structures of the cortactin SH3 domain bound to a proline-rich ligand have been reported, both involved irregular features. A complex between SH3 and the proline-rich binding motif of the interaction partner AMAP1 contained the binding motif KRPPPPPPG, suggestive of class I motif (rather than the expected class II for cortactin) and revealed an unusual 1:2 peptide:SH3 stoichiometry and a non-crystallographic twofold symmetry [50]. Another complex is a heterotrimeric 1:1:1 SH3:ligand:lysozyme assembly, with the peptide ligand derived from Arg kinase. Here lysozyme, a contaminant from SH3-purification, co-crystallized with the SH3:Arg complex, while highly purified cortactin failed to crystallize with the peptide [51]. The ligand sequence, SSVVPYLPRLPIL, contained the class VIII motif (PxxPxxP), but lacked the flanking basic residue. Generally, low affinity of SH3:PRR complexes (usually within the 10-100 μM range) hinders crystallization attempts. This underscores the importance of our first NMR-based structure of a cortactin-SH3:PRR peptide complex, circumventing crystallization artifacts and offering a native view of the complex.

SH3 domains are ubiquitous protein interaction modules that have evolved to engage proline-rich motifs found in various biological regulation cascades. Since they exhibit specific binding profiles critical for proper biological function and mediated by critical residues within the conserved scaffold despite their common fold and pre-formed binding site, it is important to identify specific molecular determinants contributing to the cortactin:Pyk2 signaling event. Our structure identifies two such PRR2 elements, the double arch P^208^xxP^211^xxP^214^ consensus motif that straddles the cortactin Y541 and W525, and the interaction of residues P214 and Y216 with a second cortactin hydrophobic surface formed by residues I521, W525, and L536. The latter - the novel of the two – is further strengthened by the K215-D522 salt-bridge and was confirmed in the course of our mutagenesis analysis. This non-canonical hydrophobic binding surface, also termed the specificity pocket [24], specificity zone [33], or surface II [52], is an essential feature of the cortactin SH3 domain and enhances cortactin affinity to ligands other than Pyk2-PRR2 (Sokolik and Chill, unpublished results). It coincides with the surface confirmed for several SH3 domains to interact with extended motifs and non-consensus ligands [33, 53], as seen from an alignment of SH3 domains [36]. Our structure emphasizes the importance of the hydrophobic packing of residues in the extended Pyk2-derived motif with surface II, in excellent agreement with the definition of the cortactin SH3 consensus sequence as +PPΨPXKP**<u>XWL</u>** [34].

Despite the high significance of cortactin for invadopodia-mediated tumor cell invasiveness and subsequent cancer metastasis, the design of a specific cortactin inhibitor has been challenging. Research to date has focused on disruption of the upstream signaling pathway EGFR-Src-Arg-cortactin or inhibition of cortactin protein-protein interactions. Indeed, inhibition of EGFR or Src suppresses invadopodia-mediated invasiveness of head-and-neck squamous cell carcinoma cell lines [54], while knockdown or selective inhibition of Arg kinase block *in vivo* breast cancer metastasis [5, 8]. Also, inhibition of the interaction between the ARF-GAP AMAP1 and cortactin by an AMAP1-derived peptide blocked tumor metastasis and angiogenesis [50, 55]. Our data establish the molecular and cellular mechanisms by which Pyk2-PRR2 inhibits cortactin and suggest that this interface is a favorable target for preventing invadopodia-mediated metastasis of breast tumor cells.

Peptides derived from endogenous protein sequences are a promising starting point for drug development, with inherent advantages including widespread utilization as receptor ligands in signaling cascades and cost-effective synthesis and purification. While they also pose challenges, such as proteolysis, rapid renal clearance, low oral bioavailability, and membrane permeability, these can be effectively overcome using peptidomimetic compounds involving D- or non-proteogenic amino acids, various cyclization approaches, and tailored delivery methods. A recent investigation found multi-layered affinity-encoding attributes in κ-helices [27], suggesting that our Pyk2-PRR2-derived peptide may have independent importance as a potential inhibitor. Furthermore, our structural understanding of their interaction with cortactin can guide the optimization of inhibition for such ligands. One logical approach would be a gain in interaction enthalpy by addition of positive charge, targeting SH3 negatively charged surfaces, or by increasing the hydrophobicity at the surface II interface. Alternatively, the entropy loss accompanying binding could be mitigated by pre-stabilization of the polyproline helix with additional proline residues or non-native amino acids. Both approaches rely upon our structural results to highlight variable peptide positions at which such modifications can occur without impairing the SH3:PRR2 interaction.

As only cancer cells make invadopodia, disrupting the mechanisms of their formation and activation by invadopodia-directed metastasis inhibitors should have minimal side effects compared to anti-cancer cytotoxic drugs currently in use. Moreover, since such inhibitors will not induce direct selective pressure on the survival of cancer cells but only inhibit invasion signaling pathways, resistance to such inhibitors is not predicted to develop. Peptide inhibitors targeting the Pyk2:cortactin interaction may confer high specificity and reduce side effects in metastatic breast cancer patients. Although a PRR2-Pyk2 peptide may not effectively block the growth of primary breast tumors. such an inhibitor should effectively inhibit the metastatic spread of highly invasive tumors when used in combination with standard cytotoxic treatments such as chemotherapy, radiotherapy, or immunotherapy.

## MATERIALS AND METHODS

### Peptides

Pyk2-PRR2 (^202^TAFQEPPPKPSRPKYRPPP^220^), Pyk2-PRR3 (^251^EFTGPPQKPPRLGAQS^266^), Pyk2 super-mutant (TAFQEP^**208**^**A**PK^**211**^**A**S^**213**^**Q**^**214**^**A**^**215**^**Q**YRPPP; a mutant of Pyk2-PRR2; P208A, P211A, R213Q, P214A, K215Q), scrambled control peptide (PEPYPTPAPFPKKQRPRPS), Pyk2-PRR2-Y216A mutant (TAFQEPPPKPSRPK^**216**^**A**RPPP), and Pyk2-PRR2 Δ214-220 (^202^TAFQEPPPKPSR^213^) were synthesized by GL Biochem (Shanghai, China). In all cellular biology and *in vivo* experiments, peptides contained the HIV-TAT sequence at their C-terminal (GRKKRRQRRRK) to increase their permeability into cells [46]. For invadopodia, matrix degradation, 2D motility assays, and 3D invasion assays, peptides containing the TAT sequence were conjugated to FITC at their C-termini to track their delivery into cells.

### Cell culture

MDA–MB-231 human breast adenocarcinoma cells and 4T1 mouse mammary tumor cells were obtained from the American Type Culture Collection. MDA-MB-231 were cultured in DMEM/10% FBS, and 4T1 cells were cultured in RPMI 1640/10% FBS. MDA-MB-231 cells stably expressing cortactin-TagRFP were previously described [22].

### Mouse xenograft model

All experimental procedures were conducted following the Federation of Laboratory Animal Science Association (FELSA) and were approved by the Bar-Ilan University animal use and care committee. Mouse xenograft tumors were generated by injecting 10^6^ 4T1 cells resuspended in 20% collagen I (BD Biosciences, Franklin Lakes, NJ, USA) in PBS into the lower-left mammary gland of 8-week-old BALB/c female mice. Twelve days following injection, when tumors reached the size of 100 mm^3^, mice were treated by intravenous injection with 10 mg/kg of control scrambled, Pyk2-PRR2, Pyk2-PRR3, or Pyk2 super-mutant peptides once a day for eight days.

### Spontaneous lung metastasis clonogenic assay

4T1 cells have intrinsic resistance to 6-thioguanine, which enables precise quantification of metastatic cells, even when they are disseminated at sub-microscopic levels at distant organs, upon excision and digestion of selected organs. The clonogenic assay is based on a principle by which one colony represents one metastatic cell [56]. Briefly, spontaneous lung metastasis was measured in BALB/c mice bearing orthotopic 4T1 tumors of equal size (1-1.2 cm in diameter). Lungs were excised and minced and plated in RPMI/10%FBS in the presence of 6-thioguanine to select for tumor cells only. Cells were plated in a 5% CO2 incubator at 37°C for 14 days to form 4T1 colonies. On day 14 after plating, the medium was removed from dishes, and colonies were fixed in 4% paraformaldehyde for one hour at room temperature. Colonies were then stained in 2% crystal violet solution and thoroughly washed with PBS. Plates were allowed to air dry, and colonies in each plate were quantified.

### Statistical analysis

Statistical analysis was performed using GraphPad Prism 8.0. For XTT, cortactin phosphorylation, and barbed end assays, statistical significance was calculated using two-way ANOVA followed by Bonferroni post-hoc test. For all other cellular and *in vivo* assays, one-way ANOVA followed by Tukey’s post-hoc test was used. For patient database analysis, statistical significance was calculated using log-rank and Student’s *t*-test implemented in MATLAB. Values were considered statistically significant if the P < 0.05. For all figures, *, P < 0.05; **, P < 0.01; ***, P < 0.001. Error bars represent standard errors of the mean (SEM).

Other experimental procedures, including NMR spectroscopy and structure determination, invadopodia assays, and proteogenomic analysis are described in detail in Supplementary Information.

## ACKNOWLEDGEMENTS

We wish to thank Dr. Avraham Samson, Dr. Moshe Dessau and Ms. Trishna Saha for technical assistance and advice, and Mr. Jonathan Solomon and Ms. Michal Gendler for critical reading of the manuscript. This work was funded by the Israel Cancer Research Fund (grant number 20-101-PG), the Israel Cancer Association (grant number 20210071) and the Israel Science Foundation (grant number 2142/21) (to Hava Gil-Henn), and the Israel Science Foundation (grant number 964/19) (to Jordan Chill).

## AUTHOR CONTRIBUTIONS

Conceptualization: J.H. Chill, H. Gil-Henn; formal analysis: S. Twafra, C.G. Sokolik, Tomer Meirson; funding acquisition: J.H. Chill, H. Gil-Henn; investigation: S. Twafra, C.G. Sokolik, T. Sneh, K.D. Srikanth, A. Genna; supervision: J.H. Chill, H. Gil-Henn; writing - original draft: J.H. Chill, H. Gil-Henn; writing - review and editing: S. Twafra, C.G. Sokolik, T. Sneh, K.D. Srikanth, A. Genna, J.H. Chill, H. Gil-Henn.

