## Supplementary material for "A novel Pyk2-derived peptide inhibits invadopodia-mediated breast cancer metastasis"

### SUPPLEMENTARY INFORMATION

#### Supplementary materials and methods

##### Proteogenomic analysis

Data from two cohorts of 1,080 and 825 breast invasive carcinoma samples was obtained from The Cancer Genome Atlas (TCGA, https://cancergenome.nih.gov/) [1] and analyzed with cBioPortal tools (http://www.cbioportal.org) [2] using MATLAB. The analyzed datasets contain mRNA-seq expression Z-scores (RNA-Seq V2 RSEM) and Agilent microarray mRNA Z-scores computed as the relative expression of an individual gene and tumor to the expression distribution of all samples that are diploid for the gene. Putative copy number alteration (GISTIC) [3] was collected from both cohorts. GISTIC is an algorithm that attempts to identify significantly altered regions of amplification or deletion and uses discrete copy number calls: homozygous deletion; heterozygous loss; neutral; gain; high amplification. Breast cancer samples are annotated with OS and/or DFS time and censorship status. Samples were split into high and low expressing groups based upon median mRNA expression Z-scores. Associations between Z-scores and patient survival (DFS and OS) were assessed by Kaplan-Meier time-event curves and Mantel-Haenszel hazard ratios using an implementation of Kaplan-Meier log rank testing from MATLAB Exchange. All statistical tests were two-sided. Mass-spectrometry based proteomic characterization of 102 breast cancer tumor samples [4] was obtained from the Clinical Proteomic Tumor Analysis Consortium (CPTAC) Data Portal (https://cptac-data-portal.georgetown.edu) [5] with cBioPortal using MATLAB. For each protein target, Z-scores were determined and associated P-values were calculated across all samples.

##### Distant metastasis-free survival analysis

Microarray datasets with Distant Metastasis-Free Survival (DMFS) annotation were obtained from the NCBI Gene Expression Omnibus (GEO) data repository [6] of high-throughput microarray experimental data. The meta-cohort dataset comprised of 1,650 tumor expression profiles of primary invasive breast cancer based on the Affymetrix U133 GeneChip microarray platform. Queried transcripts included 22,283 probe sets common to all microarrays in all study populations. Assembly of the datasets was performed using MATLAB (The Mathworks, Inc.) and GEO series (GSE) files were extracted via GEO accessions GSE11121, GSE25055, GSE7390, GSE25065, GSE17705, GSE12093, GSE1456, GSE5327 and GSE45255. The datasets have been retrieved with uniform normalization of probe intensities with MAS 5.0 [7] using global scaling with a trimmed mean target intensity of each array arbitrary set to 600. Cross-population batch effects were corrected using Z-score transformation [8]. The tumor profiles represent primary invasive breast tumors sampled at the time of surgical resection, annotated with DMFS time and censorship status Patient samples were split into high and low expressing groups based upon median gene expression. Associations between normalized gene expression and patient survival (DMFS) were assessed by Kaplan-Meier time-event curves and Mantel-Haenszel hazard ratios using an implementation of Kaplan-Meier log rank testing from MATLAB Exchange. All statistical tests were two-sided. The synergy index (SI) was calculated as a means of evaluating additive interaction [9, 10]. The synergy index can be interpreted as the excess risk from overexpression of both genes relative to the risk of overexpression of the genes separately. SI of 1 indicates no synergism, and an SI >1 indicates synergistic interaction between the two genes.

**Antibodies**

For immunofluorescence, anti-cortactin (ab-33333) and anti-PCNA (ab18197) were obtained from Abcam (Cambridge, UK); anti-Arp2 (N1C3) was obtained from GeneTex (Irvine, CA, USA); anti-Tks5 (SH3#4; 09-268) was obtained from EMD Millipore (Burlington, MA, USA); anti-pY421-cortactin (C0739) was obtained from Sigma-Aldrich (St. Louis, MO, USA); anti-pY402-Pyk2 (44-618G) was obtained from Thermo Fisher Scientific (Waltham, MA, USA); anti-CD31 (550274) was obtained from BD Biosciences (Franklin Lakes, NJ, USA); anti-cleaved caspase 3 (Asp175; 9961) was obtained from Cell Signaling Technology (Danvers, MA); Rhodamine-labelled phalloidin and Alexa Fluor-conjugated secondary antibodies were obtained from Molecular Probes (Thermo Fisher Scientific). For immunoblotting, anti-cortactin (clone 4F11; 05-180) was obtained from EMD Millipore; anti-β-actin (clone AC-15; A5441) was obtained from Sigma-Aldrich. Secondary antibodies (goat anti-mouse 680LT and goat anti-rabbit 800CW) were obtained from LI-COR Biosciences (Lincoln, NE, USA).

**Metastasis assay in a xenograft mouse model/tumor immunohistochemistry**

4T1 cells were injected into the mammary fat pad of 8-week-old BALB/c female mice and allowed to grow for 12 days. On day 13 following injection, mice were treated by intraperitoneal injection with scrambled control, Pyk2-PRR2, or Pyk2-PRR3 peptides once a day for eight days. Mice bearing 4T1 mammary tumors were then sacrificed, and tumors or lungs were excised, fixed in 4% paraformaldehyde overnight, washed for 1 hour in cold PBS, and dehydrated overnight in 30% sucrose. Next, tissue was embedded in OCT, and 5 μm thick cryostat sections were placed on silane-coated slides and dried at room temperature followed by permeabilization with 0.1% Triton X-100 for 15 min and blocking in 1% BSA and 1% FBS for 1 hour at room temperature. Samples were then incubated overnight at 4°C with the indicated primary antibodies, washed, and incubated with the appropriate secondary antibodies. Nuclei were counterstained with 4',6-diamino-2-phenylindole (DAPI). Tissue was imaged using an inverted laser scanning confocal microscope (Zeiss LSM780; 63x, NA 1.4, oil objective, ZEN black edition acquisition software) (Oberkochen, Germany). For all assays results were based on $n$ = 7-10 mice/tumors per group.

**Cell permeability assay**

MDA-MB-231 were plated and allowed to adhere for 4 h. Adherent cells were incubated with 0.1 or 1 μM Pyk2 peptides containing HIV-TAT sequences and conjugated to FITC for 20 min, 8 h, and 24 h. At the end of the incubation time, the cells were fixed and analyzed by fluorescence microscopy.

**XTT assay**

MDA-MB-231 cells were resuspended to reach a concentration of 5,000 cells/well and were seeded in triplicates in flat bottom 96 well microtiter plates. Following overnight incubation at 37°C in a tissue culture incubator, Pyk2-PRR2, Pyk2-PRR3, or a scrambled control peptide was added to a final concentration of 0.1, 1, 5, or 10 µM, and cells were followed every 24 h for 96 h in total. Activated XTT solution was prepared according to the manufacturer's instructions (Biological Industries, Beit Haemek, Israel) and added to the cells for 4 h. Two filters were used, the specific absorbance filter at 475 nm and the non-specific absorbance filter at 660 nm. Specific absorbance was calculated using the formula: A 475 nm (test) - A475 nm (blank) - A660 nm (test).

**2D single-cell random migration assay**

The 2D single-cell random migration assay was performed as previously described [11]. In brief, 6-well microtiter plates were coated with 10 µg/ml fibronectin and blocked with 1% denatured BSA. Peptide pre-treated cells were plated at 20,000 cells/well in DMEM/10% FBS containing peptides and allowed to adhere for 12-16 h. Plates were placed in a 37°C heated chamber, and images were collected using the IncuCyte® Zoom platform (Essen Bioscience, Ann Arbor, MI, USA; 20x, NA 0.60, air objective). Phase images were collected every one hour for a total of 12 h using the IncuCyte® acquisition software. Trajectory plots, accumulated distance (total cell path length), Euclidean distance (the shortest distance between the starting point and endpoint of migration), and velocity were calculated using the chemotaxis and migration tool (ibidi GMBH, Gräfelfing, Germany).

**3D scratch wound assay**

The 3D scratch wound assay was performed as previously described [12]. Briefly, 96 well Image Lock microtiter plates (Essen Bioscience) were coated with 100 µg/ml Matrigel (cat # 354234; Corning, New-York, NY, USA). Following overnight incubation at 37°C, MDA-MB-231 cells were plated at a final concentration of 350,000 cells/ml and allowed to adhere for 12-16 h in the presence of Pyk2-PRR or control peptides. Following wounding of the bottom Matrigel-cells layer, a top layer of Matrigel was added to a final concentration of 2 mg/ml and allowed to polymerize for 30 min at 37°C. Pyk2-PRR or control scrambled peptides were then added again in DMEM/10%FBS to a final concentration of 0.1, 1, or 5 μM, with or without the MMP inhibitor GM6001 (25μM). Plates were placed in a 37°C heated chamber, and images were collected using the IncuCyte® Zoom platform (Essen Bioscience; 20x, NA 0.60, air objective). Phase images were collected every one hour for a total of 12 h using the IncuCyte® acquisition software.

**Acceptor photobleaching FRET assay**

Acceptor photobleaching experiments were performed as previously described [11]. In brief, cortactin-TagRFP MDA-MB-231 cells were plated on Alexa 405 gelatin in a medium containing Pyk2-PRR or scrambled control peptides at 0.1, 1, and 5 μM concentration, fixed, and immunostained with pY402-Pyk2/Alexa Fluor 488-goat anti-rabbit (donor) and cortactin/Alexa Fluor 555-goat anti-mouse (acceptor). As a control, cells were stained with Tks5/Alexa Fluor 488-goat anti-rabbit (donor) and cortactin/Alexa Fluor 555-goat anti-mouse (acceptor). Co-localization of cortactin with degradation holes in Alexa 405-labeled gelatin was used to identify mature degrading invadopodia. For all FRET experiments, cells were imaged in PBS at room temperature on a laser-scanning inverted confocal microscope (Zeiss LSM780; 60x, NA 1.4, oil objective). A region of interest surrounding invadopodia was bleached using 100% 561-nm laser power (acceptor channel), and images were acquired in both the 488 nm and the 561 nm channels before and after bleaching. FRET efficiency was calculated as the $E$ = 1 - (donor pre/donor post) in background-subtracted images and was corrected for fluctuations in laser power and donor bleaching in ImageJ.

**Invadopodium precursor formation assay**

The invadopodium precursor formation assay was performed as previously described [11]. Briefly, gelatin was conjugated to Alexa 405 dye (Thermo Fisher Scientific). MatTek dishes were treated with 1 N HCl and coated with 50 μg/ml poly-L-lysine. A 0.2% gelatin solution was prepared in PBS, and a 1:20 mixture of Alexa 405-labeled gelatin/unlabeled gelatin was warmed to 37°C before addition to the poly-L-lysine coated plates. Gelatin was cross-linked with 0.01% glutaraldehyde followed by quenching with 5 mg/ml sodium borohydride. 125,000 MDA-MB-231 cells were plated on Alexa 405-labeled gelatin plates in the presence of 0.1, 1, or 5 μM of Pyk2-PRR peptides or scrambled control for 4 h. At the end of incubation, cells were fixed in 3.7% paraformaldehyde, permeabilized with 0.1% Triton X-100, blocked with 1% FBS, 1% BSA in PBS, and labeled with anti-Tks5 and anti-cortactin. Images were acquired using an inverted fluorescent microscope (Leica AF6000; 63x, NA 1.4, oil objective, Leica LAS AF acquisition software) (Wetzlar, Germany) equipped with an ORCA-Flash 4.0 V2 digital CMOS camera (Hamamatsu Photonics, Shizuoka, Japan). Invadopodium precursors were identified as Tks5- and cortactin-rich punctate structures found in the ventral plane of the cell that do not co-localize with degradation areas, whereas mature invadopodia were identified as Tks5 and cortactin-rich puncta that co-localize with degradation areas at the ventral plane of the cell.

**ECM degradation assay**

The *in vitro* matrix degradation assay was performed as previously described [11]. Briefly, MatTek dishes were treated with 2.5% gelatin/2.5% sucrose, cross-linked with 0.5% glutaraldehyde, treated with 10 μg/ml of fluorescently labeled fibronectin (Alexa 568; Invitrogen, Thermo Fischer Scientific), and then with 1 mg/ml NaBH4 in PBS. 125,000 cells were plated on the fibronectin/gelatin matrix in Pyk2-PRR or control peptides and allowed to degrade for 24 h. Cells were then fixed in 3.7% paraformaldehyde, and random fields were imaged using an inverted fluorescent microscope (Leica AF6000; 40x, NA 1.3, oil objective). ECM degradation was analyzed by quantifying the average degraded area in pixels per field using ImageJ.

**Immunofluorescence analysis of cortactin tyrosine phosphorylation at invadopodium precursors**

Twenty-four hours before each experiment, cells were plated on fibronectin/gelatin unlabeled matrix in the presence of a Pyk2-PRR or control peptides at 0.1, 1, and 5 μM concentration and allowed to attach for eight hours. Cells were then starved in DMEM/0.5% FBS for 16 h in the presence of peptides and stimulated with 2.5 nM EGF for 3 min or left un-stimulated (0 min EGF). Cells were fixed and stained with mouse phospho-specific cortactin antibody (pY421). To image invadopodia Z-plane, we focused on the ventral surface of the cell where the leading edge containing cortactin is focused. We then analyze the pixels corresponding to invadopodia dots, not the whole cell. The intensity (mean grey value (mgv) minus background) of cortactin-TagRFP and pY421-cortactin at invadopodium precursors was quantified. The pY421-cortactin/cortactin-TagRFP ratio was calculated as a measure of tyrosine-phosphorylated cortactin at cortactin-rich puncta. Data were normalized to resting cells (0 min EGF) and presented as the relative fold change in cortactin tyrosine phosphorylation.

**Barbed end formation assay**

The barbed end assay was performed using biotin-conjugated actin as previously described [11]. In brief, 24 h before each experiment, cells were plated on fibronectin/gelatin unlabeled matrix in the presence of Pyk2-PRR or control peptides at 0.1, 1, or 5 μM concentration and allowed to attach for 8 h. Cells were starved for 16 h in the presence of the peptides and then stimulated with 2.5 nM EGF and permeabilized with a permeabilization buffer (20 mM Hepes, pH 7.5, 138 mM KCl, 4 mM $MgCl_2$, 3 mM EGTA, 0.2 mg/ml saponin, 1 mM ATP, and 1% BSA) containing 0.4 μM biotin-conjugated muscle actin (AB07) (Cytoskeleton Inc., Denver, CO, USA) for 1 min at 37°C. Cells were fixed in 3.7% paraformaldehyde for 5 min, blocked in PBS containing 1% FBS, 1% BSA, and 3 μM un-labeled phalloidin (Molecular Probes, P3457), then labeled with FITC anti-biotin (200-092-211) (Jackson ImmunoResearch Laboratories, West Grove, PA, USA) to visualize barbed ends, and with rhodamine-phalloidin (R415) (Molecular Probes, Eugene, OR, USA) and Arp2 to identify regions of cells rich in invadopodia. The barbed end intensity at invadopodia-rich regions was quantified by measuring mgv at invadopodia-rich areas minus mgv of the background. Data were normalized to the control condition for each experiment.

**Expression and purification of cortactin-SH3 domain**

The DNA sequence encoding the SH3 domain of the mouse cortactin SH3 domain (Gene Bank ID U03184; nucleotides1468-1641) was sub-cloned into a pET-28a plasmid (Novagen, Merck) in frame with the cleavable N-terminal 6×His tag. Isotopically labeled cortactin-SH3 for heteronuclear NMR experiments was expressed in transformed *E. coli* BL21 cells (DE3) grown in the presence of kanamycin (50 mg/l) and chloramphenicol (34 mg/l) at 37 °C in M9 minimal medium [13]. For $^{15}N$ ($^{13}C$) labeling the medium contained 1 g/L of $^{15}NH_4Cl$ (4g/L $^{13}C_6$-glucose), and 0.25 g/l of appropriately labeled Celtone Base Powder (Cambridge Isotope Laboratories, Andover, MA) was added. For triple-labeled $^{2}H$,$^{13}C$,$^{15}N$-SH3, expression began in 100 ml of the above medium grown to an $OD_{600}$ of 0.4-0.5 and gently centrifuged. Pellets were then resuspended in 500 ml filter-sterilized $D_2O$-based $^{15}N$-M9 supplemented with $^{2}H_7$-$^{13}C_6$-glucose (2 g/l) and DCN-Isogro (0.25 g/l) to reach a final $OD_{600}$ of ~0.1. In both cases cells were then grown at 37 °C and induced with 0.5 mM isopropyl β-D-1-thiogalactopyranoside (IPTG) upon reaching $OD_{600}$~0.8, followed by overnight growth at 27 ºC. To purify the SH3 domain, cells were harvested by centrifugation, subjected to a single freeze-thaw cycle, and resuspended in lysis buffer (50 mM NaPi pH 8.0, 300 mM NaCl, 1 mM dithiothreitol (DTT), 100 ml per liter of culture) supplemented with 1 tablet EDTA-free protease inhibitor cocktail (Merck) and benzamidine (5 mM). Cells were lysed by French press, followed by addition of benzamidine (final concentration 10 mM) and phenylmethylsulfonyl fluoride (PMSF, 1 mM), and the lysate was clarified by centrifugation. The supernatant was loaded on a HisTrap HP column (GE Healthcare, Chicago, IL, USA) equilibrated with lysis buffer and the protein eluted through a gradient (Buffer B: 50 mM NaPi pH 8.0, 100 mM NaCl, 300 mM imidazole, 1 mM DTT). To the fraction eluting at ~40% B purified tobacco etch virus (TEV) protease (1:30 mol:mol) was added, and the mixture dialyzed overnight at 4 °C against dialysis buffer (50 mM NaPi at pH 8.0, 100 mM NaCl, 1 mM DTT). The dialysate was brought to a final concentration of ~5 mM imidazole, filtered, and loaded onto a HisTrap column. The collected flowthrough was dialyzed against 20 mM Tris (pH 7.5), 1 mM DTT for 3 h to reach a final concentration of 10 mM NaCl. This fraction was then loaded onto a Q HP Sepharose ion-exchange column (GE Healthcare) and eluted through a gradient at ~45% buffer B (20 mM Tris pH 7.5, 1 M NaCl, 1 mM DTT), flash-frozen and stored at -80 °C.

**Preparation of the SH3-PRR2 complex**

Purified SH3 was dialyzed against NMR sample buffer (30 mM NaPi, pH ~6.5) and concentrated to the desired concentration in an Amicon concentration tube (3 kDa MWCO) (Merck). The PRR2 peptide was dissolved in $D_2O$ to a final concentration 14-fold higher than the SH3 domain and combined with the SH3 sample, affording SH3/PRR2 complex in 7% $^2H_2O$ solution to which $NaN_3$ (0.02% v/v) was added. Final sample concentrations were 0.7-1.0 mM at a 1:1 molar ratio SH3:PRR2. For quantitative titrations, purchased peptides were dissolved in 100-300 µl NMR sample buffer, and samples at varying (SH3:PRR2 1:0-1:6, SH3 concentration constant at 0.1 mM) ratios were prepared. Fittings were performed with 5-6 measured points.

**NMR spectroscopy**

NMR measurements were conducted on a DRX700 Bruker spectrometer (Bruker BioSpin GmbH, Rheinstetten, Germany) with a cryogenic triple-resonance TCI probe head equipped with z-axis pulsed-field gradients at 16.4 T and 296-298 K. External calibration was performed with DSS dissolved in the same buffer as for the experiments. Acquisition parameters for 2D TOCSY, NOESY and COSY homonuclear experiments, 2D HSQC experiments, the triple resonance HNCACB experiment, and 3D filtered-edited NOESY experiments are detailed below. For two-dimensional homonuclear experiments (COSY, TOCSY, and NOESY) 2048 complex points were acquired in the direct dimension ($^1$H) (acquisition time: 208.9 ms) and 280-480 complex points in the indirect dimension ($^1$H) (acquisition times: 40-57.1 ms), with mixing times of 150 ms (TOCSY) or 200-300 ms (NOESY). Typical experiment times were 3-36 h. Conditions for filtered-edited NOESY were 1024 complex points in the direct dimension ($^1$H), 96-97 complex points in the indirect $^1$H dimension, and 25 complex points in the $^{13}$C dimension, affording acquisition times of 104.4 ms, 4.2-5.3 ms, and 17.1 ms, respectively, and a mixing time of 200-400 ms throughout 59-66 h. $^{15}$N,$^1$H-HSQC spectra were acquired at 1024 complex points in the $^1$H dimension and 188 complex points in the $^{15}$N dimension (acquisition times: 91.8 ms and 88.3 ms, respectively). The HNCACB experiment (90 h) was acquired at 1024 complex points in the direct $^1$H dimension and 40 (52) complex points in the indirect $^{15}$N ($^{13}$C) dimension, affording acquisition times of 104.4, 16.6, and 4.9 ms, respectively. The aromatic-TROSY $^{13}$C,$^1$H correlation spectrum (1.5 h) was acquired with 1024 complex points in the direct $^1$H dimension and 100 complex points in the indirect $^{13}$C dimension. Assignment of the unbound SH3 domain was assisted by BMRB entry 10240. Chemical shift perturbation values representing a normalized Euclidean distance $\Delta\delta$ were calculated as $((\Delta_H)^2+(0.14*\Delta_N)^2)^{½}$, where $\Delta_H$ and $\Delta_N$ are the shifts in the hydrogen and nitrogen dimensions, respectively.

**NMR affinity determination**

The $^1$H, $^{15}$N-HSQC spectrum of a 0.2 mM sample of the SH3 domain was acquired at 0:1 (black), 0.5:1, 1:1, 2:1 and 4:1 (light green) mol:mol PRR2:SH3 and 0:1 (black), 0.75:1, 1.5:1, 3:1 and 6:1 (light green) mol:mol PRR3:SH3. Peptides were unlabeled in all cases. Isotherms were fitted to the equation

$$\alpha = \frac{n + 1 + \frac{K_D}{C} \pm \sqrt{\left(n + 1 + \frac{K_D}{C}\right)^2 - 4n}}{2}$$

where α is the fraction of bound SH3, $n$ is the mol:mol equivalent ratio, $K_D$ the dissociation constant and $C$ the SH3 domain concentration (0.2 mM) maintained constant by preparing multiple samples with an identical amount of SH3 stock solution. α was calculated as $CSP/CSP_{max}$. Both $K_D$ and $CSP_{max}$ were treated as fitted parameters based on multiple measurements of CSP at various $n$ values.

**Structure calculation**

NOE crosspeaks were ranked by intensity and binned into 4 groups corresponding to distances of 2.7, 3.3, 4.0-4.5 and 5.0-5.5 Å. Dihedral angle constraints were derived from the TALOS+ server [14] from SH3 domain HA, HN, CA and CB chemical shifts. Initial structures were calculated using distance and dihedral angle constraints by simulated annealing and torsional angle dynamics as applied in the program CNS [15]. Final structures were calculated with CNS using distance, dihedral, and H-bond constraints in addition to two electrostatic constraints, SH3(E506)-PRR2(R213) and SH3(D522)-PRR2(K215) suggested by the NOESY data and derived from characteristic salt-bridge patterns in SH3 complexes. Stereochemical quality of the ensemble of final structures was evaluated using with VADAR 1.8 [16] and PROCHECK [17]. For the final ensemble of 20 structures, 100 structures were generated, and structures with lowest-energy and highest stereochemical quality were chosen by analysis with PROCHECK. Final coordinates were deposited at the Protein Data Bank under accession code 7PLL.

#### Supplementary figure legends

**Fig. S1 Pyk2-PRR peptides are permeable and do not affect cell viability.** Adherent MDA-MB-231 cells were incubated with 0.1 μM **(A)** or 1 μM **(B)** FITC labeled control or Pyk2-PRR peptides for 20 min, 8 h, and 24 h and analyzed by fluorescence microscopy. Shown are representative images from each group. Scale bar, 20 μM. **(C-F)** Viability of MDA-MB-231 cells in the presence of scrambled (black), Pyk2-PRR2 (red), Pyk2-PRR3 (blue), or non-treated (green) was measured every 24 h for 96 h total using the XTT reagent. $n = 3$ independent experiments, each experiment was performed in triplicates.

**Fig. S2 Pyk2-PRR2 affects the 2D random migration of breast cancer cells.** MDA-MB-231 cells were plated on fibronectin-coated plates in the presence of Pyk2-PRR or control scrambled peptides, placed at 37°C in Incucyte, and imaged every hour for 24 h. **(A, E, I)** Trajectory plots demonstrating random cell motility in 2D at 0.1 μM (A), 1 μM (E), and 5 μM (I). **(B-D, F-H, J-L)** Quantification of motility parameters: accumulated distance (total path length) (B, F, J), Euclidean distance (the shortest distance between the starting point and the endpoint of migration) (C, G, K), and velocity (D, H, L). $n$ = 49-53 (scrambled), $n$ = 45-58 (Pyk2-PRR2), $n$ = 43-51 (Pyk2-PRR3) cells from three independent experiments. * $P < 0.05$, ** $P < 0.01$, *** $P < 0.001$, as determined by one-way ANOVA followed by Tukey's post-hoc test. Error bars represent SEM.

**Fig. S3 Chemical shift perturbations of the SH3:PRR2 binding interface.** $^{1}$H,$^{15}$N-HSQC of $^{15}$N-labeled SH3 domain exhibiting chemical shift perturbation (CSP) effects induced by the addition of the PRR2 peptide (1:1 mol:mol). Cross peaks are annotated by residue, and black arrows highlight large CSPs.

**Fig. S4 Assignment of the $H_N$ and $H_\alpha$ regions of the SH3/PRR2 NOESY spectrum. (A)** Intra-PRR2 cross-peaks between amide and aliphatic sidechain protons, enabling the assignment of PRR2 chemical shifts. Spectrum was acquired for a sample of $^{2}H$-labeled SH3 in complex with unlabeled PRR2 peptide at 296 K and 16.4 T. Red annotations show the amide PRR2 proton. Black annotations indicate the aliphatic sidechain proton involved. **(B)** Intra-PRR2 cross-peaks between $H_{\alpha}$ and aliphatic sidechain protons, enabling the assignment of PRR2 chemical shifts. This is particularly important for a Pro-rich peptide since proline residues lack the key $H_{N}$-proton used for assignment. Spectrum was acquired for a sample of $^{2}H$-labeled SH3 in complex with unlabeled PRR2 peptide at 296 K and 16.4 T. Red annotations show the amide PRR2 proton. Black annotations indicate the aliphatic side chain proton involved.

**Supplementary movies**

**Supplementary Movie 1. 3D motility of MDA-MB-231 cells treated with control (scrambled) peptide.** MDA-MB-231 were embedded in Matrigel and allowed to invade into an artificially made wound. Invading cells were imaged using IncuCyte® every hour for 12 h (20x/NA 0.6, air objective).

**Supplementary Movie 2. 3D motility of MDA-MB-231 cells treated with Pyk2-PRR2 peptide.** MDA-MB-231 cells were embedded in Matrigel and allowed to invade into an artificially made wound in the presence of 5 μM Pyk2-PRR2 peptide. Invading cells were imaged using IncuCyte® every hour for 12 h (20x/NA 0.6, air objective).

**Supplementary Movie 3. 3D motility of MDA-MB-231 cells treated with Pyk2-PRR3 peptide.** MDA-MB-231 were embedded in Matrigel and allowed to invade into an artificially made wound in the presence of 5 μM Pyk2-PRR3 peptide. Invading cells were imaged using IncuCyte® every hour for 12 h (20x/NA 0.6, air objective).

**Supplementary Table 1. Structural statistics for the SH3-PRR2 complex.**

<table>
<tr><td colspan="2">NMR distance restraints</td></tr>
<tr><td>Total inter-residue constraints</td><td>811</td></tr>
<tr><td>Sequential (|i-j|=1)</td><td>258</td></tr>
<tr><td>Medium range (1< |i-j| < 5)</td><td>76</td></tr>
<tr><td>Long-range (|i-j|>5)</td><td>477</td></tr>
<tr><td>Total intra-SH3 constraints</td><td>645</td></tr>
<tr><td>Total intra-peptide constraints</td><td>47</td></tr>
<tr><td>Total inter-molecular constraints</td><td>119</td></tr>
<tr><td colspan="2">NOE violations, Å</td></tr>
<tr><td>Maximum single violation</td><td>0.5</td></tr>
<tr><td>Rmsd of NOE violations</td><td>0.23</td></tr>
<tr><td colspan="2">Dihedral angle restraints</td></tr>
<tr><td>TALOS-derived dihedral restraints</td><td>106</td></tr>
<tr><td colspan="2">Dihedral angle violations, °</td></tr>
<tr><td>Maximum single violation</td><td>5</td></tr>
<tr><td>Rmsd of dihedral violation</td><td>0.24</td></tr>
<tr><td colspan="2">Deviation from idealized geometry</td></tr>
<tr><td>Bond lengths, Å</td><td>0.002</td></tr>
<tr><td>Bond angles, °</td><td>0.38</td></tr>
<tr><td>Improper angles, °</td><td>0.28</td></tr>
<tr><td colspan="2">Mean rmsd values, Å</td></tr>
<tr><td>Backbone (SH3, PRR2 207-215)</td><td>0.21</td></tr>
<tr><td>Heavy atoms (SH3, PRR2 207-215)</td><td>0.83</td></tr>
</table>

**Figure S1**

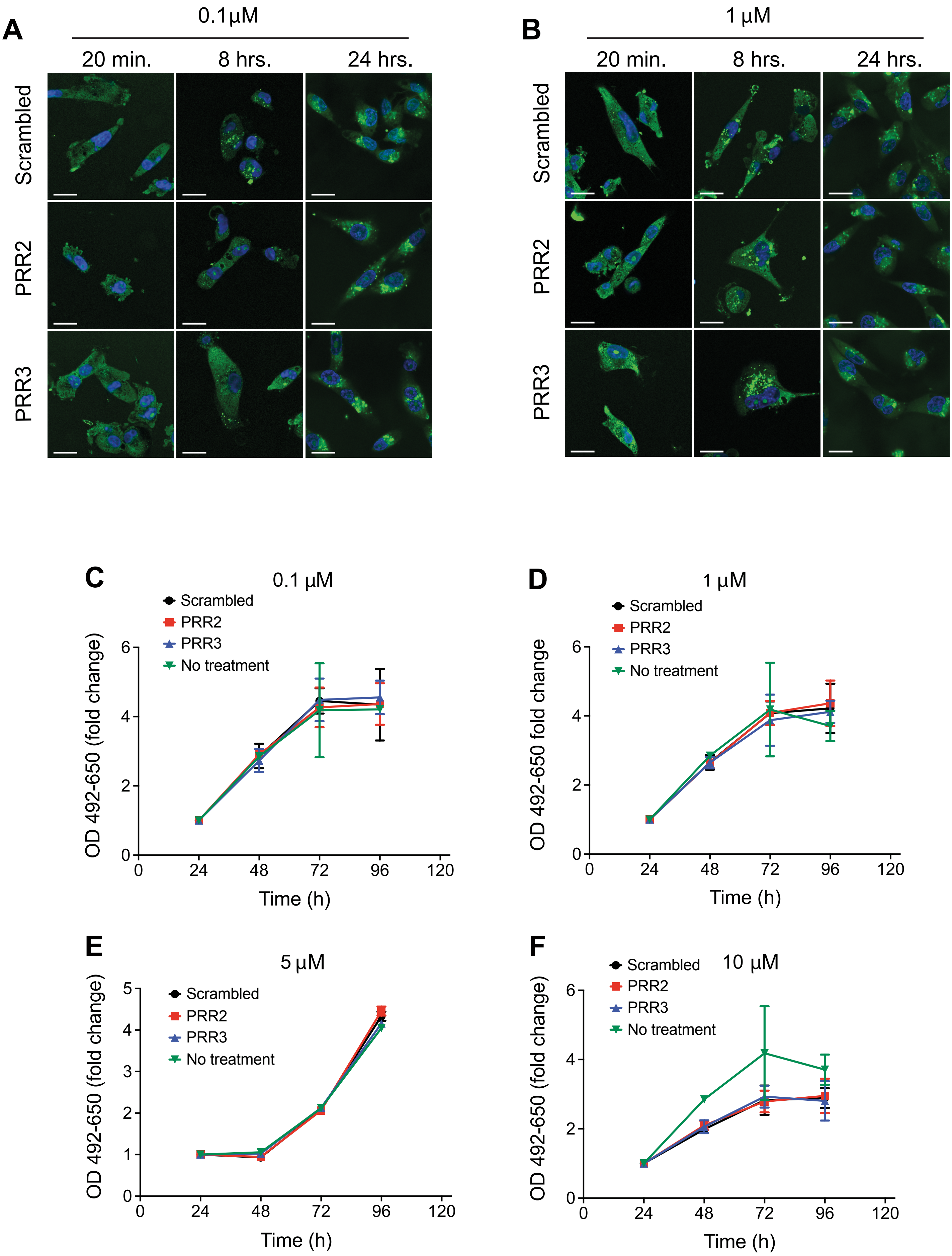

A
0.1µM
20 min.
8 hrs.
24 hrs.
Scrambled
PRR2
PRR3
B
1 µM
20 min.
8 hrs.
24 hrs.
Scrambled
PRR2
PRR3
C
0.1 µM
Scrambled
PRR2
PRR3
No treatment
OD 492-650 (fold change)
Time (h)
D
1 µM
Scrambled
PRR2
PRR3
No treatment
OD 492-650 fold change
Time (h)
E
5 µM
Scrambled
PRR2
PRR3
No treatment
OD 492-650 (fold change)
Time (h)
F
10 µM
Scrambled
PRR2
PRR3
No treatment
OD 492-650 (fold change)
Time (h)

### Figure S2

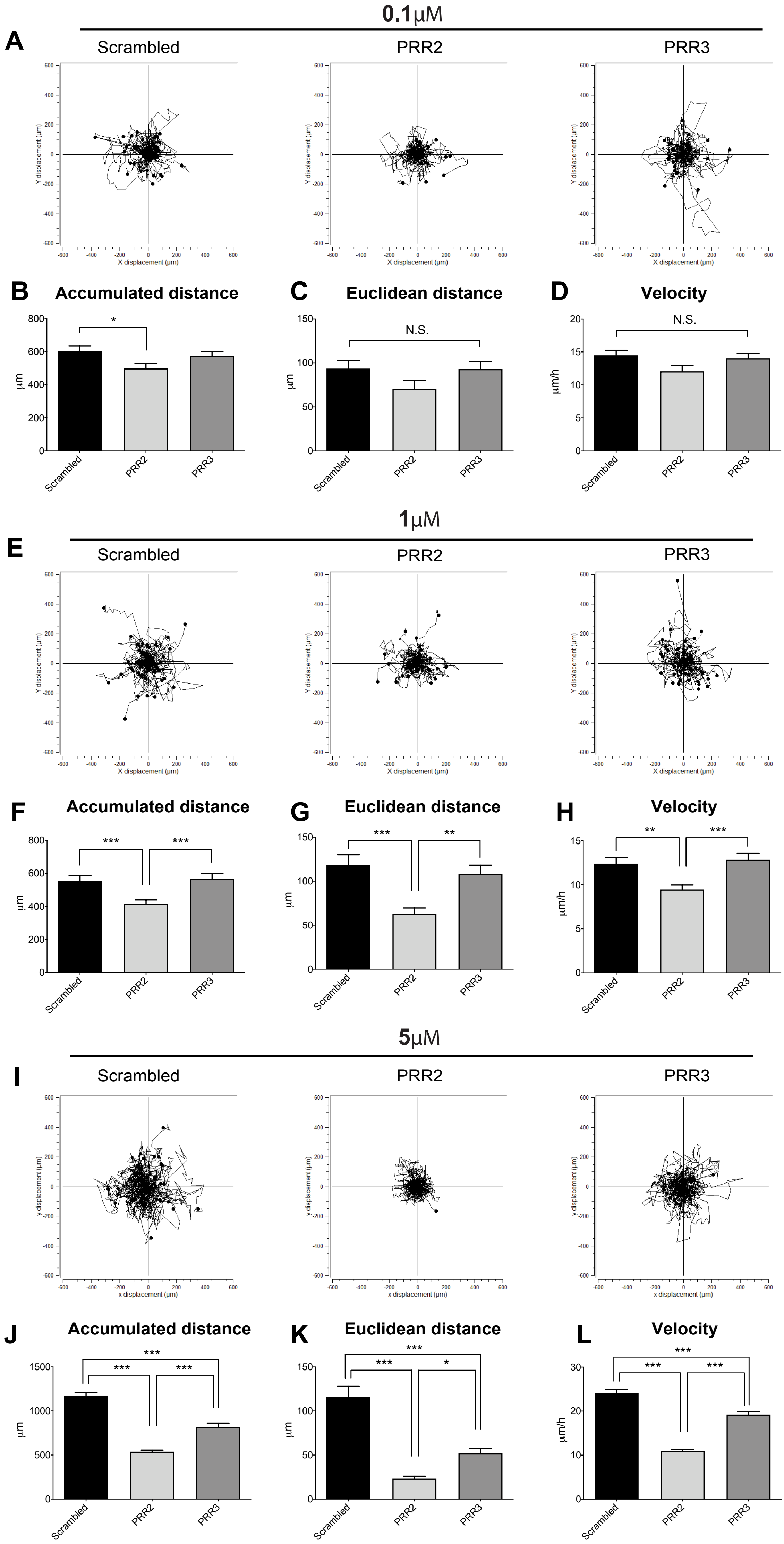
0.1μM
A
Scrambled
PRR2
PRR3
Y displacement (μm)
X displacement (μm)
B Accumulated distance
C Euclidean distance
D Velocity
μm
μm/h
N.S.
Scrambled
PRR2
PRR3
1μM
E
F Accumulated distance
G Euclidean distance
H Velocity
5μM
I
y displacement (μm)
x displacement (μm)
J Accumulated distance
K Euclidean distance
L Velocity

### Figure S3

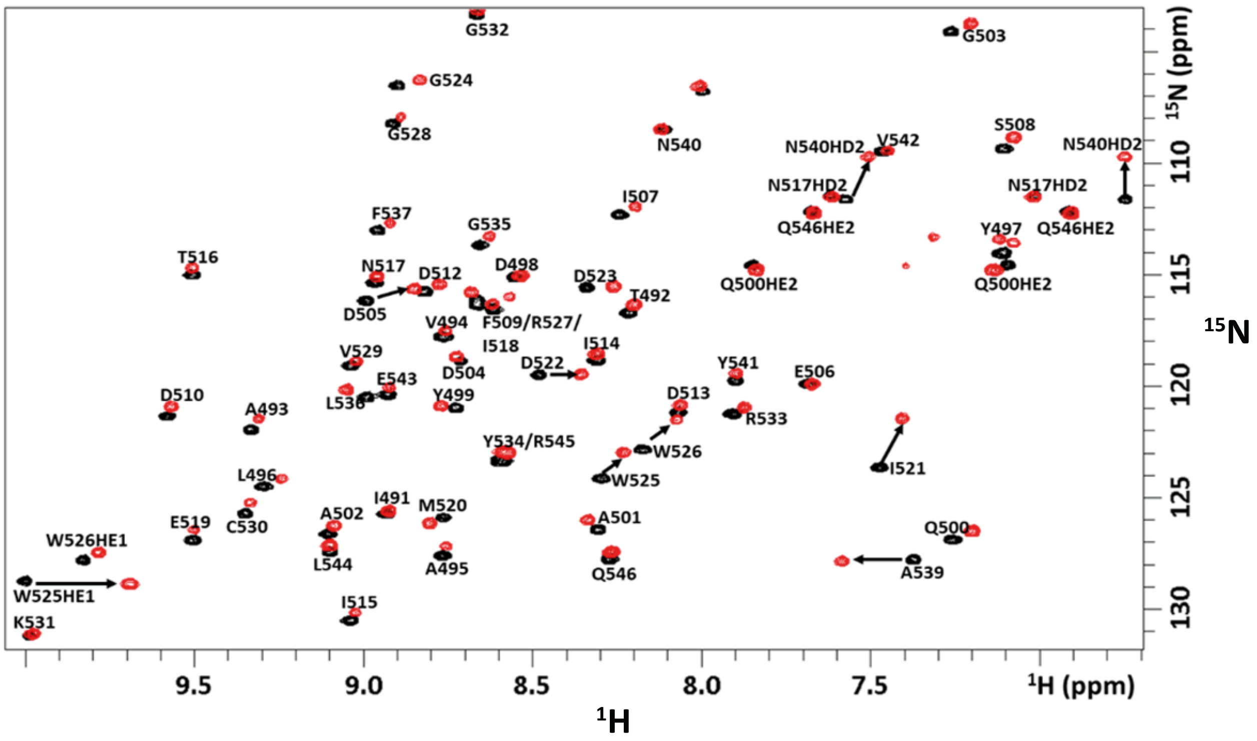

G532
G503
G524
G528
N540
S508
V542
N540HD2
N540HD2
I507
N517HD2
N517HD2
F537
G535
Q546HE2
Y497
Q546HE2
T516
N517
D512
D498
D523
T492
Q500HE2
Q500HE2
D505
V494
F509/R527/
I518
I514
V529
D504
D522
Y541
E506
E543
D510
L536
Y499
D513
A493
R533
Y534/R545
W526
I521
L496
W525
I491
M520
A502
E519
C530
A501
Q500
W526HE1
L544
A495
Q546
A539
W525HE1
I515
K531
9.5
9.0
8.5
8.0
7.5
$^{1}$H (ppm)
$^{1}$H
110
115
120
125
130
$^{15}$N (ppm)
$^{15}$N

### Figure S4

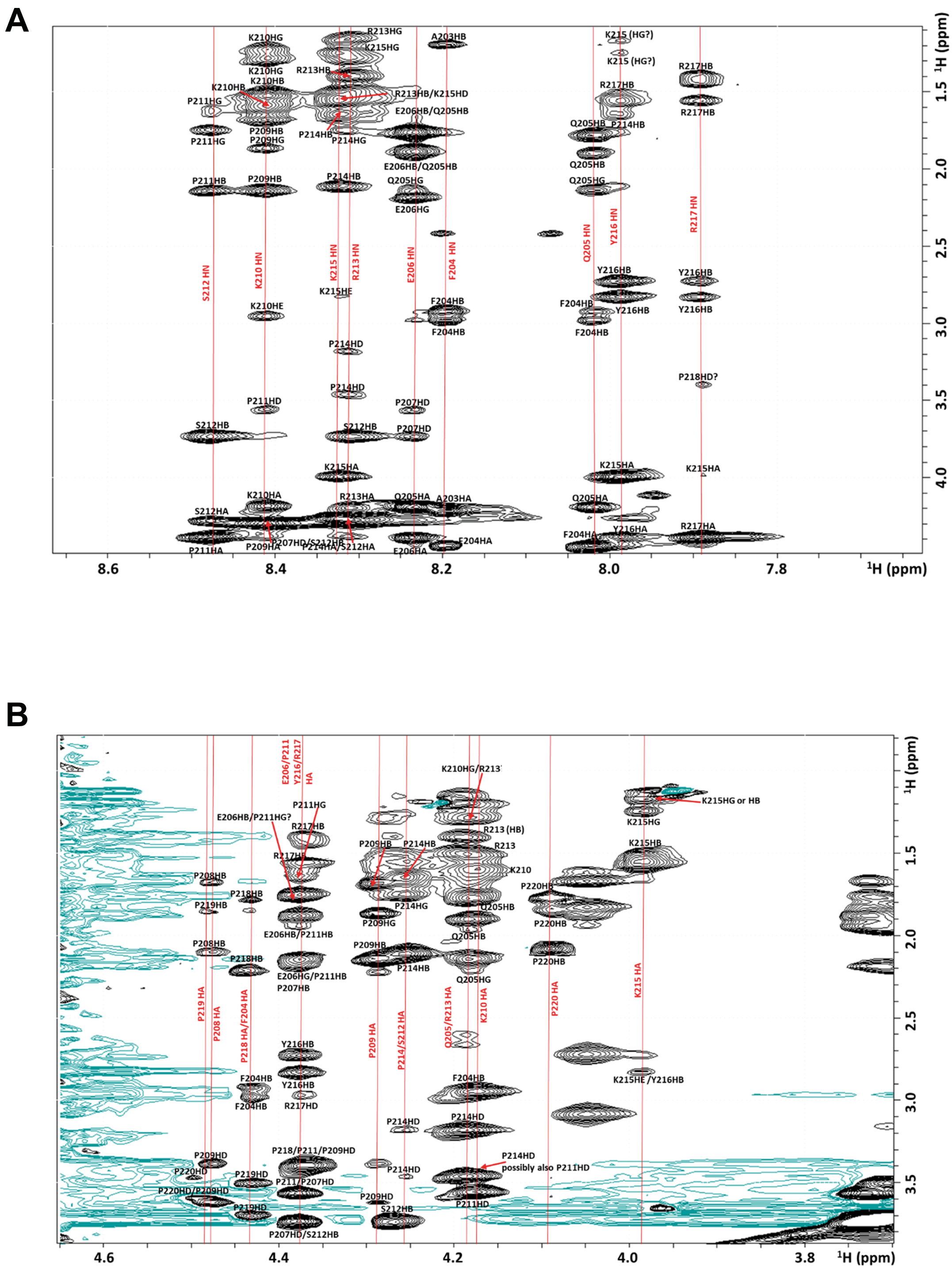
A
B
¹H (ppm)
8.6
8.4
8.2
8.0
7.8
1.5
2.0
2.5
3.0
3.5
4.0
4.6
4.4
4.2
4.0
3.8
S212 HN
K210 HN
K215 HN
R213 HN
E206 HN
F204 HN
Q205 HN
Y216 HN
R217 HN
K210HG
R213HG
K215HG
A203HB
K215 (HG?)
R213HB
R213HB/K215HD
R217HB
K210HB
P211HG
E206HB/Q205HB
P209HB
P209HG
P214HB
P214HG
Q205HB
P211HB
Q205HG
E206HG
Y216HB
K215HE
K210HE
F204HB
P214HD
P218HD?
P211HD
P207HD
S212HB
K215HA
K210HA
R213HA
Q205HA
A203HA
S212HA
Y216HA
R217HA
F204HA
P211HA
P209HA
P207HD/S212HB
P214HA/S212HA
E206HA
E206/P211
Y216/R217
HA
K210HG/R213
K215HG or HB
E206HB/P211HG?
R213 (HB)
R213
K210
P208HB
P218HB
P219HB
P220HB
E206HB/P211HB
E206HG/P211HB
P207HB
P220HB
P219 HA
P208 HA
P218 HA/F204 HA
P209 HA
P214/S212 HA
Q205/R213 HA
K210 HA
P220 HA
K215 HA
K215HE /Y216HB
R217HD
P218/P211/P209HD
P214HD
possibly also P211HD
P209HD
P220HD
P219HD
P211/P207HD
P220HD/P209HD
P211HD
P207HD/S212HB
